# Neuronal responses in mouse inferior colliculus correlate with behavioral detection of amplitude modulated sound

**DOI:** 10.1101/2021.11.02.466979

**Authors:** Maurits M. van den Berg, Esmée Busscher, J. Gerard G. Borst, Aaron B. Wong

## Abstract

Amplitude modulation (AM) is a common feature of natural sounds, including speech and animal vocalizations. Here, we used operant conditioning and *in vivo* electrophysiology to determine the AM detection threshold of mice as well as its underlying neuronal encoding. Mice were trained in a Go-NoGo task to detect the transition to AM within a noise stimulus designed to prevent the use of spectral side-bands or a change in intensity as alternative cues. Our results indicate that mice, in comparison with other species, detect high modulation frequencies up to 512 Hz well, but show much poorer performance at low frequencies. Our *in vivo* multielectrode recordings in the inferior colliculus (IC) of both anesthetized and awake mice revealed a few single units with remarkable phase-locking ability to 512 Hz modulation, but not sufficient to explain the good behavioral detection at that frequency. Using a model of the population response that combined dimensionality reduction with threshold detection, we reproduced the general band-pass characteristics of behavioral detection based on a subset of neurons showing the largest firing rate change (both increase and decrease) in response to AM, suggesting that these neurons are instrumental in the behavioral detection of AM stimuli by the mice.

**New and Noteworthy:** The amplitude of natural sounds, including speech and animal vocalizations, often shows characteristic modulations. We examined the relationship between neuronal responses in the mouse inferior colliculus and the behavioral detection of amplitude modulation in sound, and modelled how the former can give rise to the latter. Our model suggests that behavioral detection can be well explained by the activity of a subset of neurons showing the largest firing rate changes in response to AM.

## 1. Introduction

A major goal of neuroscience is to understand how sensory stimuli are encoded in the firing patterns of sensory neurons and how these patterns guide behavior. This is typically studied in animals by comparing neuronal responses with behavioral detection of a suitable stimulus. This comparison of neurometric and psychometric performance has yielded much information about the way relevant sensory features are encoded at different stages of the sensory system, but also about the contribution of other parts of the sensorimotor loop, which include non-sensory factors such as attention, motivation, bias, decision making and the motor response.

In these sensory detection tasks individual neurons may outperform the animal (Carney et al., 2014; Johnson et al., 2012; Kettner et al., 1985; Malone et al., 2010; Newsome et al., 1989). If many neurons individually outperform the animal, their individual impact on behavior would be expected to be very small. However, this does not seem to be the case, as stimulation of even a single neuron may already impact sensory detection (Tanke et al., 2018). What might account for this apparent discrepancy between the ability of individual sensory neurons to encode a stimulus or impact behavior and the comparatively poor performance of animals in sensory detection tasks? A possibility is that the activity of different sensory neurons is heavily correlated, as these so-called noise correlations would limit the information these neurons can carry (Azeredo da Silveira et al., 2021). However, noise correlations are often small (Bartolo et al., 2020; Renart et al., 2010), as also indicated by the observation that neuronal population responses can be two orders of magnitude more sensitive than the animal’s performance, indicating that there are few or no limitations due to noise correlations within the population responses in the sensory precision that can be attained (Stringer et al., 2021). The gap between neurometric and psychometric performance might then be filled by non-sensory factors (de Lafuente et al., 2006; Goris et al., 2017; Guo et al., 2014; Kettner et al., 1985; Stringer et al., 2021), but we argue here that sensory performance in some behavioral tasks may also be easily overestimated, for example because the behavioral task may include multiple types of stimuli, making the task more difficult.

Amplitude modulations (AM) forms an important cue in communication sounds such as speech, but also in the identification of environmental sounds, segregating sound sources during auditory scene analysis, pitch perception or musical perception (Joris et al., 2004; Rees et al., 2005). One way or another, the firing of each auditory neuron is modulated by sound amplitude. AM tuning is typically characterized by presenting sinusoidally amplitude modulation (SAM) of a carrier sound at different frequencies to measure how well neurons phase-lock to the modulation frequency, yielding the temporal modulation transfer function (tMTF), or how vigorously they fire in response to different modulation frequencies, yielding the rate modulation transfer function (rMTF). The tMTF changes noticeably between the cochlear nucleus and the inferior colliculus (IC). The tuning becomes sharper, and the ability of IC neurons to phase-lock to high modulation frequencies is reduced (Joris et al., 2004). Further along the pathway in the auditory cortex, the upper limit of modulation frequency to which neurons phase-lock further decreases (Joris et al., 2004), which has led to the suggestion that for behavioral detection AM sounds are encoded by a rate code rather than a temporal code (Bendor et al., 2007; Dong et al., 2011; Lu et al., 2004; Niwa et al., 2012b). However, temporal codes may represent sounds for both pure AM and natural vocalizations with large AM components more accurately (Henry et al., 2016; Schnupp et al., 2006; Walker et al., 2008; Wang et al., 2007). Even though there are many studies that have studied either the behavioral detection of AM or the electrophysiological encoding of AM (Joris et al., 2004; Rees et al., 2005), the question which mechanisms for encoding AM support behavioral sensitivity is therefore still far from settled.

Here, we performed multielectrode recordings in the IC of both anesthetized and awake mice, and compare SAM responses of single units with the behavioral performance of mice in a SAM detection task to investigate how neuronal responses in the IC may contribute to AM detection. We found that despite the large diversity and the complex tuning of many IC neurons to AM noise, a small fraction of the units explains most of the task-relevant variance in the neuronal activity. By taking the relation between modulation depth and response latency into account, we could provide an adequate description for the behavioral performance based on the neural responses. Moreover, a comparison of neuronal responses in anesthetized and awake animals indicated that the latter approximates the behavioral responses better. Our data thus provide a partial explanation for the gap that exists between neurometric and psychometric performance.

## 2. Materials and Methods

### 2.1. Animals

Wildtype B6CBAF1/JRj mice of both sexes were obtained from Janvier (behavior: n = 8 [3 females]; acute electrophysiology recordings: n = 18 [9 females; 4 from behavioral training]; awake electrophysiology recordings: n = 4 [2 females]). During the experimental period, the mice were housed with 2 to 4 mice per cage, enriched with a running wheel, an acrylic house and nest building material. Ethical approval was granted prior to the start of the experiments from the national authority (Centrale Commissie Dierproeven, The Hague, The Netherlands; IRN number: 2019-020) as required by Dutch law, and all experiments were performed according to institutional, national, and European Union guidelines and legislation as overseen by the Animal Welfare Board of the Erasmus MC.

### 2.2. Surgery

For mice used in behavioral experiments and awake electrophysiology recordings, a titanium head plate was placed on the skull of the animals for head-fixation during the experiments. The animals were anesthetised through respiratory delivery of isoflurane and maintained at surgical level of anesthesia, assessed through the hind limb withdrawal reflex. A heating pad with rectal feedback probe (40-90-8C; FHC, Bowdoinham, ME, USA) was used to maintain body core temperature at 36-37 °C. Eye ointment (Duratears^®^; Alcon Nederland, Gorinchem, The Netherlands) was used to keep the eyes moist during surgery. Boluses of buprenorphine (0.05 mg/kg; Temgesic^®^, Merck Sharp & Dohme, Inc., Kenilworth, NJ, USA) and carprofen (5 mg/kg; Rimadyl^®^, Zoetis, Capelle a/d IJssel, The Netherlands) were injected subcutaneously at the beginning of surgery. The skin overlying the IC was incised. Lidocaine (Xylocaine^®^ 10%; AstraZeneca, Zoetermeer, The Netherlands) was applied before removing the periosteum and cleaning the skull. After etching the bone surface with phosphoric acid gel (Etch Rite™; Pulpdent Corporation, Watertown, MA, USA), the titanium head plate was glued to the dorsal surface of the skull using dental adhesive (OptiBond™ FL; Kerr Italia S.r.l., Scafati, SA, Italy) and further secured with dental composite (Charisma^®^; Heraeus Kulzer GmbH, Hanau, Germany).

For acute anesthetized experiments, prior to headplate attachment, anesthesia was induced with ketamine (10%, Alfasan, Woerden, The Netherlands) and xylazine (Sedazine^®^, ASTfarma, Oudewater, The Netherlands) mixture (130 mg.kg^-1^ and 13 mg.kg^-1^, respectively) by IP injection. An intraperitoneal cannula (Butterfly Winged Infusion Set, 25G x ¾”, NIPRO EUROPE N.V., Zaventem, Belgium) was placed to allow for a maintenance solution of ketamine/xylazine to be injected. This was done with a syringe pump (Fusion 200, Chemyx, Stafford, TX, USA) providing a continuous rate infusion of between 35 - 45 mg/kg/h of ketamine and 1.4 – 1.8 mg/kg/h of xylazine. Adequacy of anesthetic depth was evaluated by toe-pinch, and the infusion rate was adjusted when needed. Headplate attachment was performed as described above. A craniotomy was performed overlying the left IC for recording. In acute experiments a second craniotomy was performed overlying the frontal lobe for the ground screw. For mice used in repeated awake recordings, a silver wire was tucked underneath the skull through the craniotomy for grounding. To protect the craniotomy and minimize shift of the brain, the craniotomy of the acute mouse was covered with 2% agarose dissolved in rat Ringer’s solution (148 mM NaCl, 5.4 mM KCl, 1 mM MgCl_2_, 1.8 mM CaCl_2_, 5 mM HEPES), while those of the awake recordings were covered in Dura-Gel (Cambridge Neurotech Ltd., Cambridge, United Kingdom).

### 2.3. Amplitude-modulated sound stimulus

Sound stimuli were generated in MATLAB R2019a (The MathWorks, Natick, MA, USA) and played back via a TDT System3 setup (a combination of RX6 processor, PA5 attenuator and ED1 electrostatic speaker drivers, or a RZ6 processor) driving two EC1 electrostatic speakers (Tucker Davis Technologies, Alachua, FL, USA). Sound stimuli were presented in open field bilaterally during behavioral and awake electrophysiology experiments, and unilaterally to the right ear in the anesthetized experiments. Sound intensities were calibrated using a condenser microphone (ACO pacific Type 7017; ACO Pacific, Inc., Belmont, CA, USA) connected to a calibrated pre-amplifier and placed at the position of the pinnae.

The base stimulus consisted of a broad-band pink (1/f) noise between 2 and 48 kHz (behavior) or between 2 and 64 kHz (electrophysiology). The broadband stimulus allowed the concurrent stimulation of neurons with a wide range of characteristic frequencies (CFs). Each trial started with an unmodulated noise with a 10 ms cosine squared ramp. After a certain duration of time, the sound switched into a sinusoidally modulated version of the base stimulus according to:

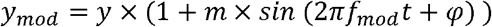

where y and y_mod_ are the base and amplitude-modulated (AM) stimuli, respectively; m is the modulation depth between 0 (no modulation) and 1 (full modulation); f_mod_ is the modulation frequency in Hz; t is time in s; and *φ* is an offset in phase to accommodate differences in instantaneous intensity (see below). In the behavioral test the animals were trained to detect the transition into amplitude modulation.

Several measures were taken to prevent the animal from detecting possible non-AM cues (Figure S1). First of all, amplitude modulation alters the power spectrum of the noise by the introduction of a pair of sidebands to each of its frequency components. We denote the collective modulation sidebands by the term “sideband spectrum.” The unmodulated part of each stimulus was generated by first creating another instance of the carrier noise (Figure S1A), calculating the sideband spectrum required to produce the desired amplitude modulation of the trial (Figure S1B-C), followed by randomization of the phase of these sidebands in the spectral domain (Figure S1D) and finally adding this randomized sideband back to the original carrier noise (Figure S1E). This procedure eliminated any spectral and intensity differences between the modulated and unmodulated parts of the noise stimulus (Figure S1F-H). Secondly, to avoid an abrupt transition, the unmodulated and amplitude-modulated noise were cross-faded with a 20-ms window following a cosine and a sine function, respectively, between 0 and π/2 (Figure S1I-L). The Pythagorean identity sin^2^θ + cos^2^θ = 1 ensured that the combined instantaneous power of the two uncorrelated noises remained constant. Thirdly, the amplitude modulation started at the phase at which the instantaneous value of the sine function matched the average power of the unmodulated noise: 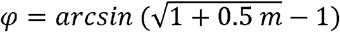, where m is the modulation depth (Figure S1I-J).

### 2.4. Behavioral training

The animals were motivated through water control. They received water only during habituation and training sessions during the week and ad libitum water during the weekends (Friday evening to Sunday afternoon). Each mouse received one session on Monday and two sessions per day for the rest of the week. Before the first training session, mice were habituated for two weeks to head-fixation in the setup with gradually increasing durations. During training sessions, animals were head-fixed and placed in a sound isolated booth between two speakers and in front of a metal lick port for dispensing water rewards. Licking behavior was measured by means of a change in capacitance of the lick port connected to an Arduino Nano utilizing the CapacitiveSensor library (playground.arduino.cc/Main/CapacitiveSensor/). Mice were trained to suppress their licking behavior after the onset of unmodulated (UM) noise until the transition to amplitude modulated noise (16 Hz, full modulation depth, 60 dB SPL), upon which licking was rewarded with a water drop. The duration of the inter-trial intervals (silent) and the delay to AM transition were randomized between 2-3 s and 0.5-3.5 s, respectively. If the animal licked during the UM period, an additional timeout delayed the start of the AM transition. We call this timeout the “no-lick period”, which was randomized per trial between 4-7 s at the testing stage. The same delay strategy was used for the silent period between trials. Additionally, unrewarded catch trials with a transition into unmodulated noise were introduced to gauge the false alarm rate. The stimuli were subsequently expanded to include other modulation frequencies, depths and sound intensities. For each session, stimuli were presented in blocks, in which all possible stimuli were presented in randomized order. Each animal completed between 13-49 trials per stimulus (mean: 28.8 ± 11.9 trials/stim). The number of catch trials was twice as high as for the other trials in each block to allow a more robust estimation of the false alarm rate.

The animals were tested with the method of constant stimuli after stable performance was reached. Modulation frequencies ranged from 4-1024 Hz in octave steps; modulation depths were 0.06, 0.125, 0.25, 0.5 and 1. We reported hit rates and false alarm rates in Figure 3A using a response window within 750 ms after the onset of the AM transition. For each AM stimulus, an ROC analysis was performed to compare its response latencies distribution to that of catch trials with the corresponding intensity. The sensitivity measure *d’* was then calculated from the AUC obtained from ROC analysis using the formula 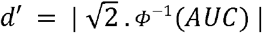 where *Φ*^-1^ represents the inverse of the standard normal cumulative distribution function (norminv in MATLAB). This method of calculating *d’* was chosen over the classical formula *Φ*^-1^(*hit rate*) − *Φ*^-1^(*false alarm rate*) [often written as z(H)-z(F)](Macmillan et al., 2004) to detect differences in response latency distributions that may not be detected using an arbitrarily chosen response window. Furthermore, the AUC method can be extended to physiological data to facilitate comparisons. The method we employed gave more conservative threshold estimates for our dataset than the classical method for calculating *d’* (data not shown). To avoid infinities in d’ for cases with perfect separation, we applied the limit *Φ*^-1^(1 − 1/2*N*_*trials*_) − *Φ*^-1^(1/2*N*_*trials*_) (Macmillan et al., 2004). For neuronal recordings (*N*_*trials*_ = 20) this amounted to a maximum *d’* of 3.92.

### 2.5. Auditory Brainstem Response (ABR)

The ABR was used to obtain hearing level thresholds of behavioral animals prior to head plate attachment surgery. Mice were anesthetized with a mixture of ketamine/xylazine (100/10 mg/kg) i.p. and placed 3.5 cm in front of an electrostatic speaker in a sound-attenuated booth. Bilateral needle electrodes were placed subdermally overlying the bulla, while the active electrode was placed at the vertex and a ground electrode placed approximately over the sacrum. Tone pips of 1 ms were presented at 4, 8, 16 and 32⍰kHz at intensities ranging from 0 to 70⍰dB sound pressure level (SPL; re 20 μPa) at 5-dB resolution. Tone pips were generated by an RZ6 multi-I/O processor, while recordings were collected on a preamplifier (Medusa4Z, Tucker Davis Technologies, Alachua, FL, USA), all controlled by BioSigRZ (Tucker Davis Technologies, Alachua, FL, USA) software. At each sound intensity tested, a minimum of 500 brainstem evoked responses with artifacts below 30⍰μV were averaged. The minimum threshold was defined as the lowest intensity at which a reproducible waveform could be identified. After the recording was completed, atipamezole (0.5 mg/kg) (Antisedan^®^, Zoetis, Capelle a/d IJssel, The Netherlands) was injected i.p. to aid in the recovery from anesthesia. All 8 animals had normal hearing thresholds (Figure S2; mean ± s.d. 4 kHz: 42.5 ± 4.29 dB; 8 kHz: 18.8 ± 2.31 dB; 16 kHz: 11.9 ± 5.30 dB; 32 kHz: 43.8 ± 7.44 dB).

### 2.6. Electrophysiological recordings

Silicon probe recordings of neuronal responses in the mouse IC to matching AM stimuli were performed both under anesthesia and in awake animals.

A multichannel silicon electrode (64 channel, single linear shank, 1250 μm, H3, Cambridge Neurotech Ltd., Cambridge, United Kingdom) was coated in DiO (3,3’-Dioctadecyloxacarbocyanine perchlorate, catalog number: D275, Invitrogen, ThermoFisher Scientific, Waltham, MA, USA) or DiI (1,1’-dioctadecyl-3,3,3’,3’-tetramethylindocarbocyanine perchlorate; catalog number: D282, Invitrogen) saturated in 97% ethanol, and inserted under direct visual inspection during online monitoring of the channels for sound-evoked spikes. The electrode was connected to an acquisition board (RHD USB-interface board, part C3100, Intan Technologies, Los Angeles, CA, United States), after A/D conversion at the headstage amplifier (RHD 64-channel headstage, part C3315, Intan Technologies) and recorded at a sampling frequency of 30 kHz using the supplied recording GUI (RHD data acquisition GUI, Intan Technologies). A TTL pulse from the RZ6 indicating the onset and offset of each stimulus was recorded on a digital channel of the Intan interface board for subsequent alignment of extracted spike times relative to the stimuli.

Pure-tone responses were recorded in FRA sets (frequency: 2-64 kHz in ¼ octave steps; intensity: 0-60 dB SPL in 5 dB steps; duration: 100 ms; intertrial interval: 200 ms; repetitions: 20, presented in pseudorandom order).

AM recording sets contained amplitude modulated noise stimuli as presented in the behavioral paradigm, except they had a fixed UM duration of 1 s before transitioning into AM, which then played for another 1 s. The intertrial interval (ITI) was set to a minimum of 500 ms. AM parameters were: modulation frequency (16, 64, 128 & 512 Hz), modulation depth (0, 0.06, 0.125, 0.25, 0.5 & 1) and intensity (45 & 60 dB SPL). Trials were presented in pseudorandomized order.

After completion of the recordings, transcardial perfusion under pentobarbital anesthesia was performed with 4% paraformaldehyde (PFA) in 0.12 M phosphate buffer. The brain was dissected out and postfixed in 4% PFA for 30 minutes. After storage in 10% sucrose, the brain was embedded in gelatin, sectioned on a freezing microtome and stained for DAPI. Images of slices (Axio Imager M2, Zeiss, Oberkochen, Germany) were used to reconstruct the location of the probe path in the IC, and aligned to the Allen Brain Atlas Common Coordinate Framework v3 (CCFv3) using Matlab code adapted from SHARP-Track (Shamash et al., 2018). Using the coordinates of the tract and the location of single units along the linear shank of the electrode, the location of individual extracted units within the brain was determined.

Single-unit activity was obtained with spike sorting using Kilosort2 (Pachitariu et al., 2016) and manual curation of the sorted results using Phy2 (https://github.com/cortex-lab/phy). Single units were selected based on good signal-to-noise ratio in the waveform with the “template amplitude” easily separable from background multiunit activities, visual inspection of the consistency of spike waveforms across the recording and adherence to a refractory period of 1 ms by inspection of the auto-correlogram.

### 2.7. Pure-tone response analysis

To map the tuning of single units to pure tones, the frequency autocorrelation areas (FACA) (Geis et al., 2011) were calculated from the kernel spike density (KSD) of the evoked spike rates. From this tuning curve, the characteristic frequency (CF, the frequency with the lowest intensity to produce a significant response) and the minimum threshold (the lowest intensity to evoke a significant response) were calculated.

### 2.8. AM analysis

Temporal coding was quantified by a measure of neural synchronization to the envelope of AM, phase-projected vector strength (VS_PP_), as described in Yin et al. (2011). In contrast to traditional vector strength, VS_PP_ tolerates lower firing rates better. Both VS_PP_ and average firing rate were calculated from the spikes occurring in the time window between the transition from UM to AM and the end of AM.

One challenge in comparing inter-unit performance using the firing rate was the large variations in firing rates between units. Additionally, in order to compare the performance of AM encoding between the temporal and rate modalities, they would need to be calculated on the same scale. To address these challenges, the calculated parameters were converted to a performance score in d’ scale using ROC analysis. For both VS and firing rate, this was done by comparing trial-by-trial measures of VS_PP_ or firing rate between modulated and unmodulated stimuli. AUC was calculated from the ROC curves, and subsequently converted to d’ (d’_temporal_ and d’_rate_, respectively), as described in 2.4 Behavioral training. By assigning a cutoff of 1 to the d’ curve, a threshold was determined for each modulation frequency through linear interpolation in dB scale.

The temporal and rate best modulation frequency (tBMF and rBMF) of a unit was determined as the modulation frequency with the lowest d’_temporal_- or d’_rate_-derived threshold, respectively. Units with identical thresholds for two or more modulation frequencies (temporal: 3/211 units) and those with no detectable threshold at any modulation frequencies (temporal: 21/211 units; rate: 10/211 units) were excluded from further analysis on BMF. Chi-squared tests were used to assess differences in BMF distributions among subregions of the IC. Where overall significance was found (p<0.05) –pairwise Chi-squared tests were performed between IC subregion, with the Holm-Bonferroni correction for multiple comparisons.

### 2.9. Principal component analysis of response differences

We used principal component analysis (PCA) to investigate how the animals might make use of the neuronal data to detect AM in the behavioral task. Since animals were trained to distinguish between UM and AM irrespective of the modulation frequency, we first calculated the difference between the averaged neuronal response to AM across all frequencies and the averaged response to UM.

The spike times of all neurons were converted into peristimulus time histograms (PSTHs) with a 3.9 ms bin size (r(t); Figure 9A-B) and averaged across the 20 repetitions of each stimulus. The small bin size was chosen in order to allow cycle-to-cycle variation and minimize bias towards rate coding. We then calculated a response difference between the average response to full modulation stimulus (*m* = 1) across all modulation frequencies and that to the unmodulated control stimulus (Δr(t); Figure 9C). This difference was combined across neurons into a T x N matrix where N = 112 neurons and T = 768 time points (3.9 ms sampling between -0.5 to 2.5 seconds relative to stimulus onset) which was used as the input for the principal component analysis (PCA). PCA was performed using the Matlab pca function with the data and the default singular value decomposition algorithm. The PCA found the principal components that best explains the difference between fully modulated and unmodulated responses. To test the robustness of the results, we systematically evaluated the effect of parameter choices for the PCA (Suppl. Note 1 and Figure S4). Each resulting principal component (PC) contains a series of N weights (*w*_*i*_), each relating to one input neuron *i*. Summing the products between all *w*_*i*_ and their respective *Δr*_*i*_*(t)* recreates the PC. Similarly, we can map (“project”) the raw population activity r(t) to the PC by multiplying the respective weight to the firing rate of each neuron: 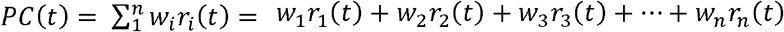 This projection operation is performed for the model below and for data shown in Figures 9E and 12.

### 2.10. Simulation of responses from neural data

We simulated how an animal would make use of this neuronal information to perform our behavioral task. Single trial spike times for each neuron were converted into a kernel spike density (KSD) function using a 2 ms Gaussian kernel and a sample interval of 0.5 ms. The combined response of all neurons was then projected into PC1 as described above. Assuming the animals were detecting a change from the unmodulated noise, the average PC1 value in the 500-ms window preceding the AM transition was taken as baseline and was subtracted from the trace. We then checked the time at which the baseline-subtracted PC1 crossed a certain “detection level” value and added an extra 100-200 ms non-decision time to get a simulated response latency, at a maximum of 1 s (stimulus duration). We systematically tested different detection level values (−200 to 200 in 2 unit steps; range of actual PC1 values: [-239.8, 444.6]) and non-decision times (100 to 200 ms in 10 ms steps) and compared them to response latencies collected from behavioral experiments. For each stimulus, empirical cumulative distribution functions of simulated response latencies were calculated from 0 to 1 s in 20 ms bins (Figure 12B), and compared to that of the behavioral data using their root-mean-square difference (Figure 12C). To compare the model with behavior, the set of 20 simulated response latencies (one for each repetition) for each stimulus was used to calculate *d’* values and estimate detection thresholds for different modulation frequencies (Figure 12D), analogous to what was performed for the behavioral test.

## 3. Results

To test for the behavioral detection of AM, we trained a total of eight mice in a Go-NoGo paradigm, in which they had to respond to the transition from an unmodulated to an amplitude-modulated (AM) broadband noise stimulus (Figure 1A). Similar paradigms have been used in a variety of species to detect psychophysical AM modulation thresholds for different modulation frequencies (Bremen et al., 2017; Klump et al., 1991; Moody, 1994). Detection thresholds were obtained by presenting a total of nine different modulation frequencies ranging from 4 to 1024 Hz at a total of five different modulation depths and two sound intensities. The stimulus was designed to specifically test for the AM cue, while controlling for both spectral and intensity differences induced by the AM (see Figure S1 and Methods). Example responses of one mouse to a low (16 Hz) and a high (512 Hz) modulation frequency at 60 dB are shown in Figure 2. The mice learned the task well, except, as is often the case with a Go-NoGo paradigm, false positive rates during catch trials were relatively high. However, they did not differ for 45 dB (32.8 ± 13.1%; range 12.2-46.1%) and 60 dB stimuli (31.7 ± 8.9%; range 21.8-44.7%; paired t-test, p = 0.68).

**Figure 1.**
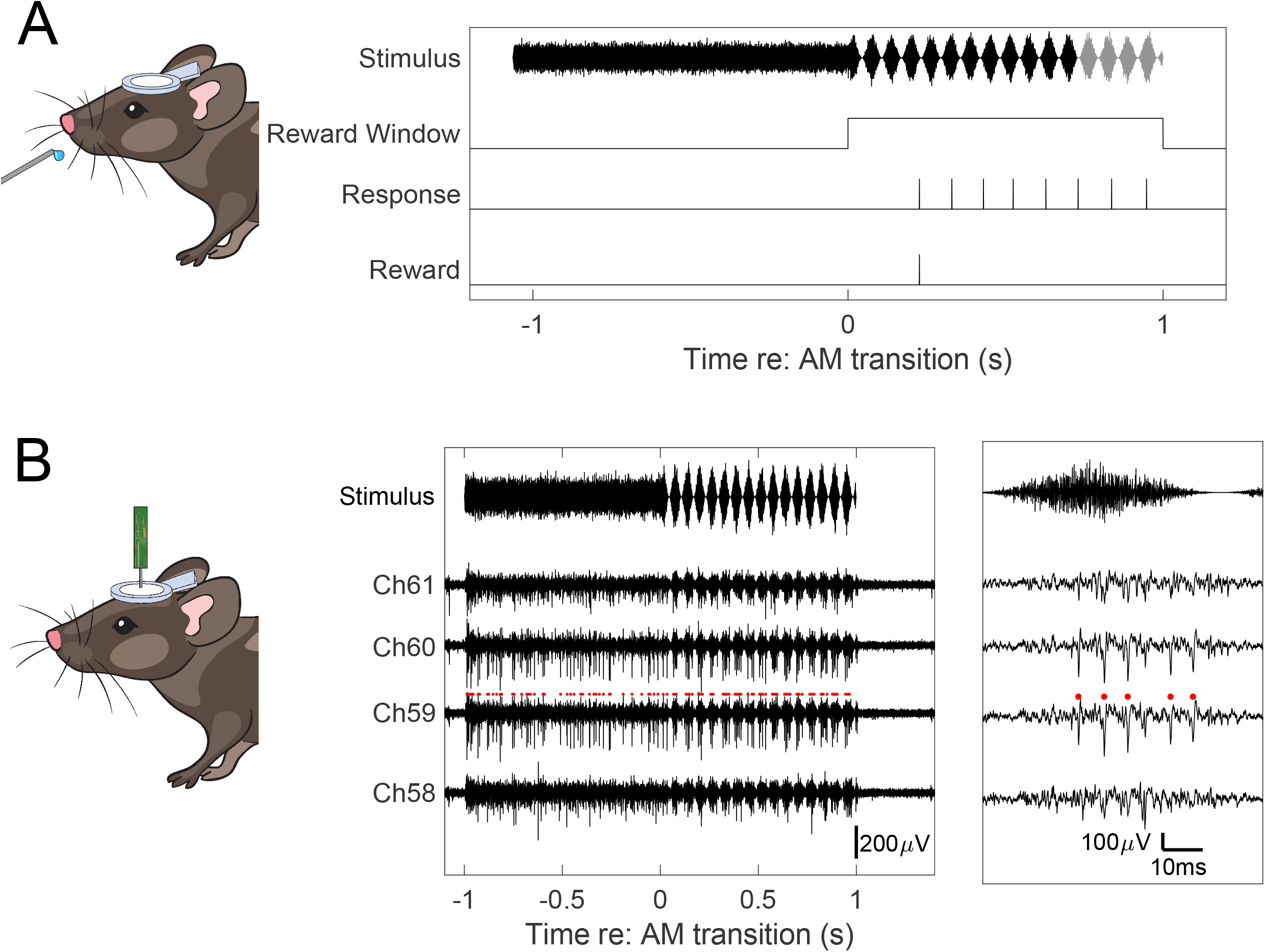
Experimental design. (A) Behavioral trials consisted of unmodulated (UM) noise stimuli of random 0.5-3 s duration that transitioned to amplitude-modulated (AM) noise stimuli with matched intensity and spectral properties. During the UM period, any lick from the mouse delayed the start of the AM transition until it did not lick for 4-7 s; the first lick within the 1 s reward window, which started at the transition to AM, was rewarded with a water drop. End of the trial was 500 ms after the response (black stimulus trace) or 1 s after its start (end of grey stimulus), whichever came first. (B) Electrophysiology trial structure. The UM and AM periods were both fixed at 1 s. Example traces from four adjacent channels is shown, and a segment of approximately one modulation period shown in the insert on the right. Red markers: spikes from a single unit.

**Figure 2.**
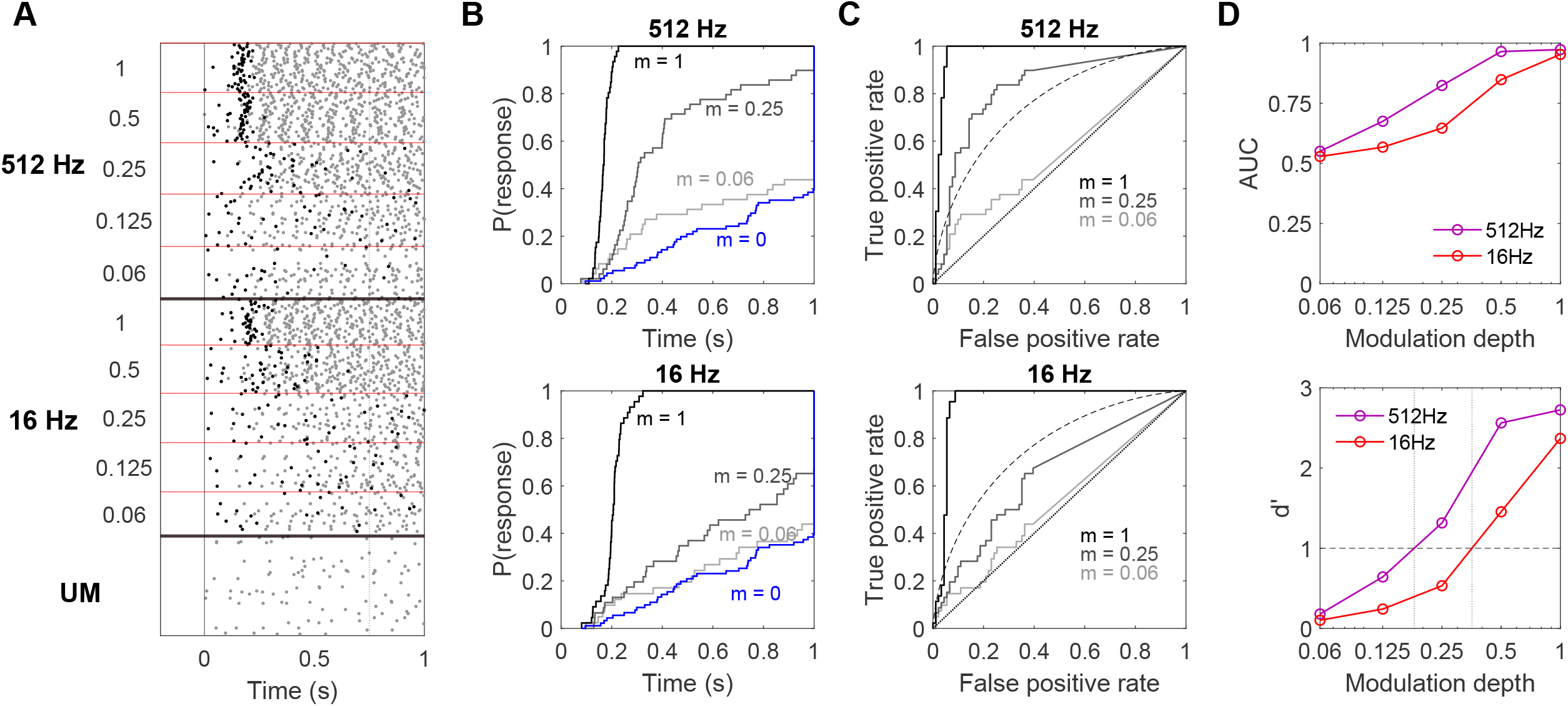
Behavioral detection of a low and a high frequency AM stimulus. (A) Dot raster of lick responses to AM transitions for an example mouse at a low and a high modulation frequency; Responses at 512 and 16 Hz modulation frequency were sorted from top to bottom for decreasing modulation depths. Black dots represent the rewarded licks; grey dots the other licks. The transition to AM occurred at t = 0 s (solid vertical line). Note that due to the “no-lick period” rule (see 2.4 Behavioral training) a transition will not be initiated unless the animal has withheld licking for a 4-7 s period, which explains the absence of licks immediately before t = 0. The dotted vertical line marks the end of the response window for calculation of hit rates in Figure 3. UM is the unrewarded catch trial condition. (B) Cumulative distribution of response latencies after the AM transition for 512 Hz (top) and 16 Hz (bottom) at various modulation depths m. Response latencies shorter than 75 ms were considered spontaneous responses and were excluded from analysis. (C) Receiver-Operator-Characteristics (ROC) analysis for response latency, plotting cumulative probability of a response to AM versus UM stimuli as the criterion (response window) changes. Dashed curve is the theoretical ROC curve for Sensitivity index (d’) = 1. (D) Top: Area-under-the-curve (AUC) of the ROC curves in C plotted as a function of modulation depth. Bottom: Sensitivity index (d’) converted from AUC values according to 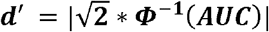, where ***Φ***^**-1**^ is the inverse cumulative distribution function of the standard normal distribution. The vertical dotted lines indicate the estimated modulation depth at which d’ = 1. All responses are from the same animal.

**Figure 3.**
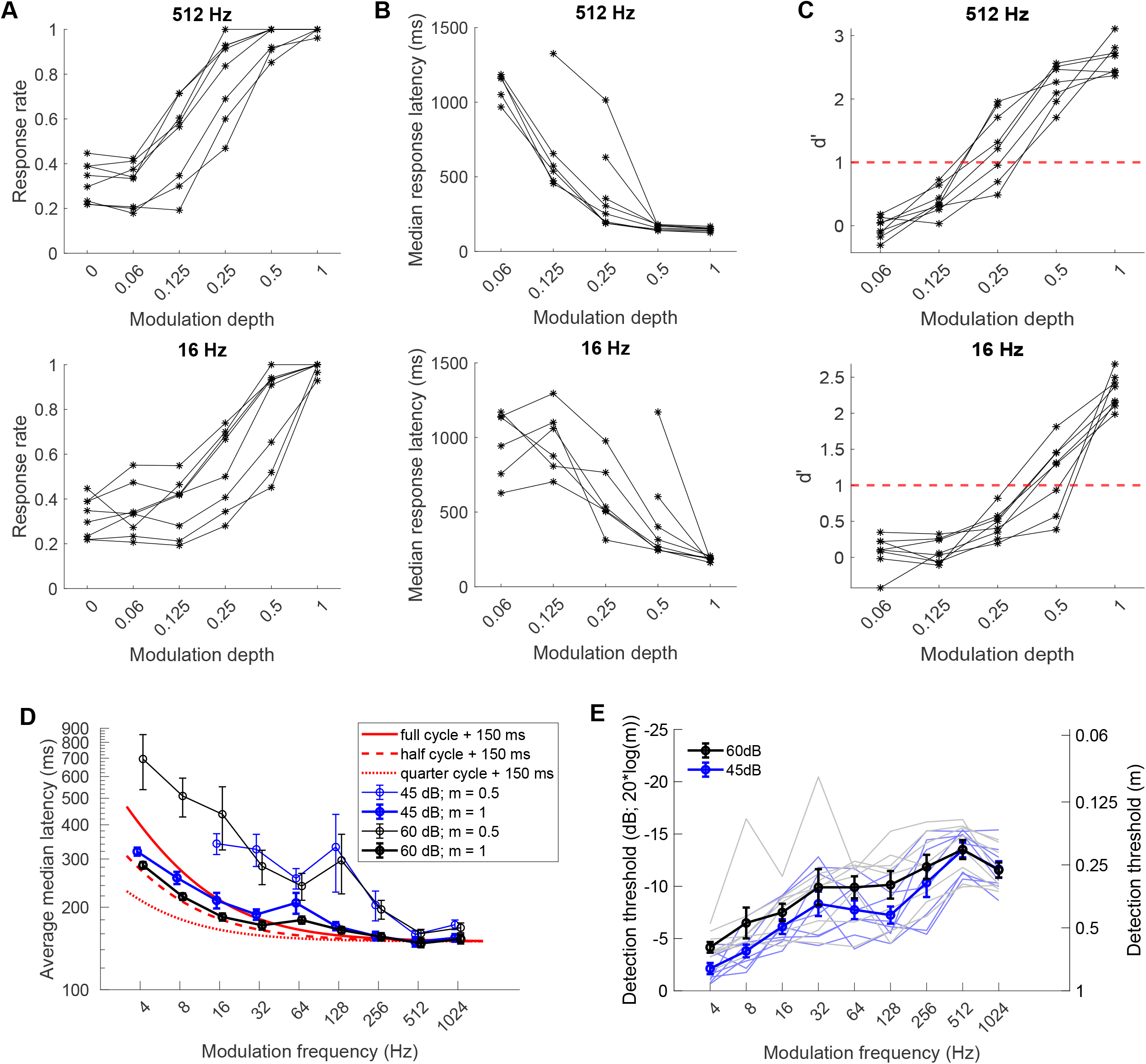
Behavioral detection thresholds at different AM frequencies show a high pass pattern. (A) Probability of a behavioral response within 750 ms after transitioning to 512 (upper) or 16 Hz (lower) AM at different modulation depths. (B) Median response latencies of animals at 512 and 16 Hz, showing faster responses at higher modulation depths. (C) Response latencies at 512 (upper) and 16 (lower) Hz were converted to d’ using ROC analysis to account for response bias. Red, horizontal dashed lines indicate d’ = 1, our definition of the detection threshold. (D) Median response latencies (mean ± s.e.m.) to AM sounds as a function of modulation frequency for 60 dB (black) and 45 dB (blue) noise. At full modulation (m = 1; thick lines), the response latencies showed high consistency across animals, and could be well approximated by a fixed delay (150 ms) plus half a modulation cycle (red dashed line). (E) Frequency- and intensity-dependence of the AM detection threshold of mice plotted as mean ± s.e.m. (thick lines). Thin lines are responses of the individual mice. Significant frequency and intensity dependence was observed. (Linear mixed effect model; frequency: p < 0.0001, intensity: p < 0.0001). However, when the effect of intensity was tested for individual modulation frequencies, none of the modulation frequencies reached statistical significance.

We used response latency as the main readout of how readily each animal detected each stimulus. To correct for response bias, response latencies during target and unmodulated catch trials were compared using ROC analysis (Figure 2B,C). Areas under the curve (AUC) were calculated and subsequently converted to the sensitivity index d’ (see Methods for rationale; Figure 2D). Higher d’ values were observed at a modulation frequency of 512 Hz than at 16 Hz, indicating that this animal was better at detecting the former. For each modulation frequency, the detection threshold was defined as the modulation depth at which d’ crossed 1 (vertical dotted lines in Figure 2D).

Figure 3 summarizes the behavioral response of all 8 animals. As expected, response probability (response window: 750 ms) for all animals increased with increasing modulation depth for a wide range of modulation frequencies (Figure 3A). Moreover, at all modulation frequencies animals responded faster to higher modulation depths (Figure 3B), most likely reflecting the certainty of the animal’s detection and subsequent response. The average median response latency to the fully modulated stimuli at 60 dB could be well approximated by adding half a period of the modulation to a fixed delay of about 150 ms (Figure 3D) suggesting that under these conditions, detection approached the physical limit at which sinusoidal amplitude modulations can be detected.

Similar to the example shown in Figure 2, we converted the response latencies of all animals into the sensitivity index d’ to estimate their detection threshold (Figure 3C). Behavioral detection of AM showed a clear improvement with modulation frequency up to 512 Hz (Figure 3E), from thresholds of about 0.5 in the lower range (8 Hz and lower) to about 0.25 in the higher range (256 Hz and higher). For instance, detection thresholds at 16 Hz were significantly higher (worse) than those at 512 Hz (Figure 3C; Linear mixed-effect model: 16 Hz vs 512 Hz, p < 0.001). The behavioral detection threshold worsened at 1024 Hz, in line with the overall low-pass filter-like high frequency limits described in many species (Carney et al., 2013; Dent et al., 2002). Overall, the mice showed a band-pass characteristics in our AM detection paradigm.

The unexpected shape of the behavioral modulation transfer function prompted us to investigate the underlying encoding mechanism. We compared the behavioral performance with neuronal responses in the IC. The IC is a key nucleus in the auditory pathway, and the first nucleus where tuning for AM responses consistently emerges (Joris et al., 2004; Rees et al., 2005).

We recorded a total of 112 well isolated single units from the IC of 18 anaesthetized mice using multielectrode silicon probes. We presented the same AM stimuli as in the behavioral task (Figure 1B, <u>S1</u>), as well as pure tones to obtain their frequency response areas (FRAs). Phase-projected vector strength (VS_PP_, (Yin et al., 2011) was used as a measure for phase-locking to the AM stimuli to study the temporal coding of modulation frequencies and the average firing rate was used to study the rate coding.

The units showed large variability in their responses to both AM stimuli and tones. In Figure 4 and 5 we illustrate the responses of two example units from the central nucleus of the IC (ICc). Figure 4 shows a unit with excellent phase-locking to AM stimuli. The unit responded to pure tones with a strong onset response followed by inhibition across a wide range of frequencies (Figure 4B). Sustained firing for the remainder of the tone duration only occurred at a narrow bandwidth centered around the CF. The frequency response area (FRA) further illustrates its sharp tuning with its CF at 16 kHz and a minimum threshold of 20 dB (Figure 4C).

**Figure 4.**
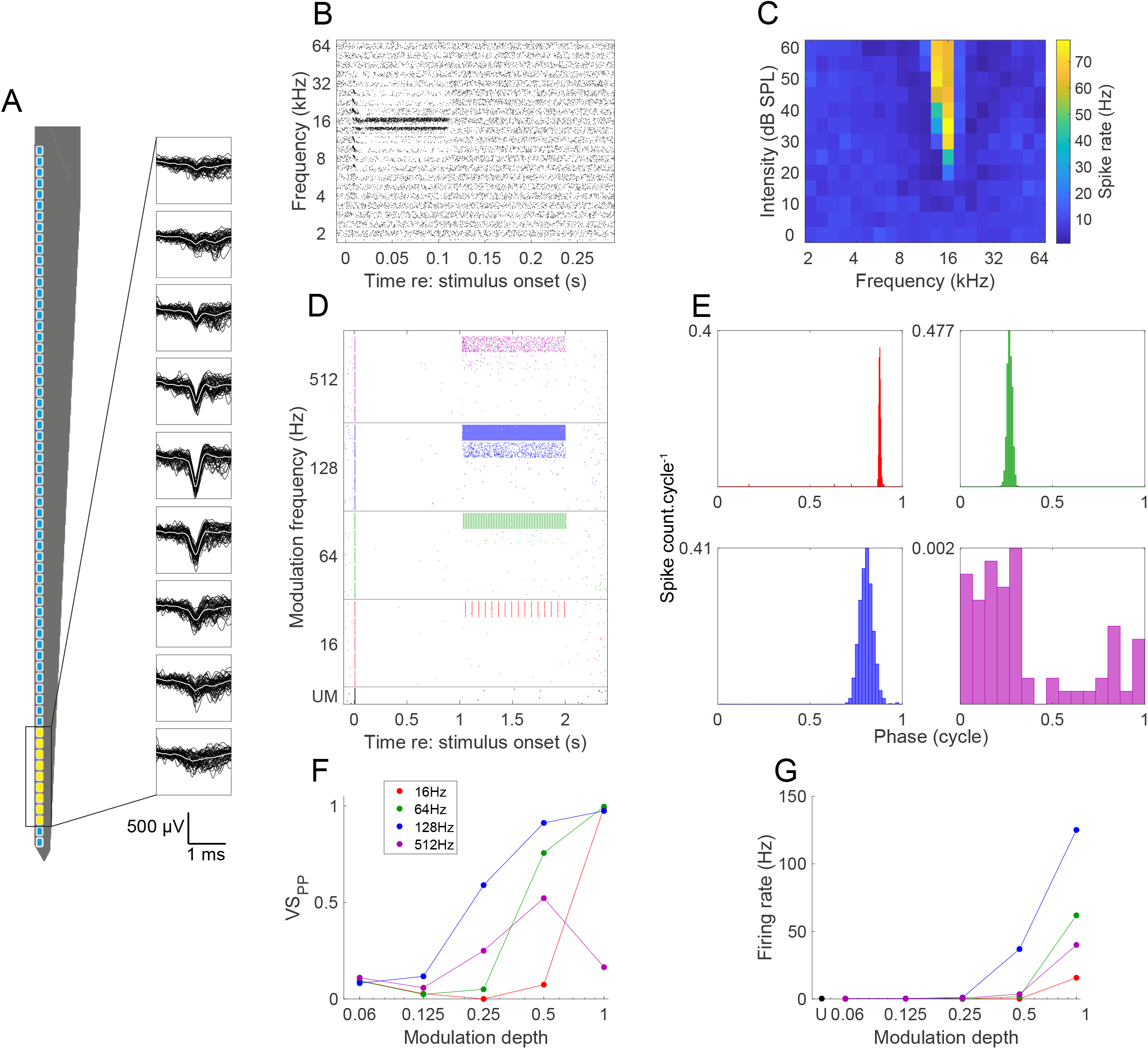
Example unit from the ICc with an onset response and relatively good phase-locking to AM stimuli. (A) Spike waveforms detected on nine adjacent channels. (B) Dot raster plot of the pure tone responses with the ordinate sorted by frequency followed by intensity. (C) Responses to 100 ms pure tones displayed as a heatmap show sharp tuning with a CF of 16 kHz and a minimum threshold of 20 dB. (D) Dot raster plot of responses to AM stimuli. Trials are organized from top to bottom by decreasing modulation depth for the different modulation frequencies. Transition from UM to AM occurred at 1 s. (E) Cycle histograms of peak phase-locking modulation depth for the same modulation frequencies as shown in D. Bin size is 0.13 ms. (F) Temporal coding of AM was quantified by calculating the phase-projected vector strength (VS_PP_) for the responses during the AM period. (G) The average firing rate during the AM period is used as a measure for rate coding to AM. All AM data shown are for a carrier noise intensity of 60 dB SPL.

**Figure 5.**
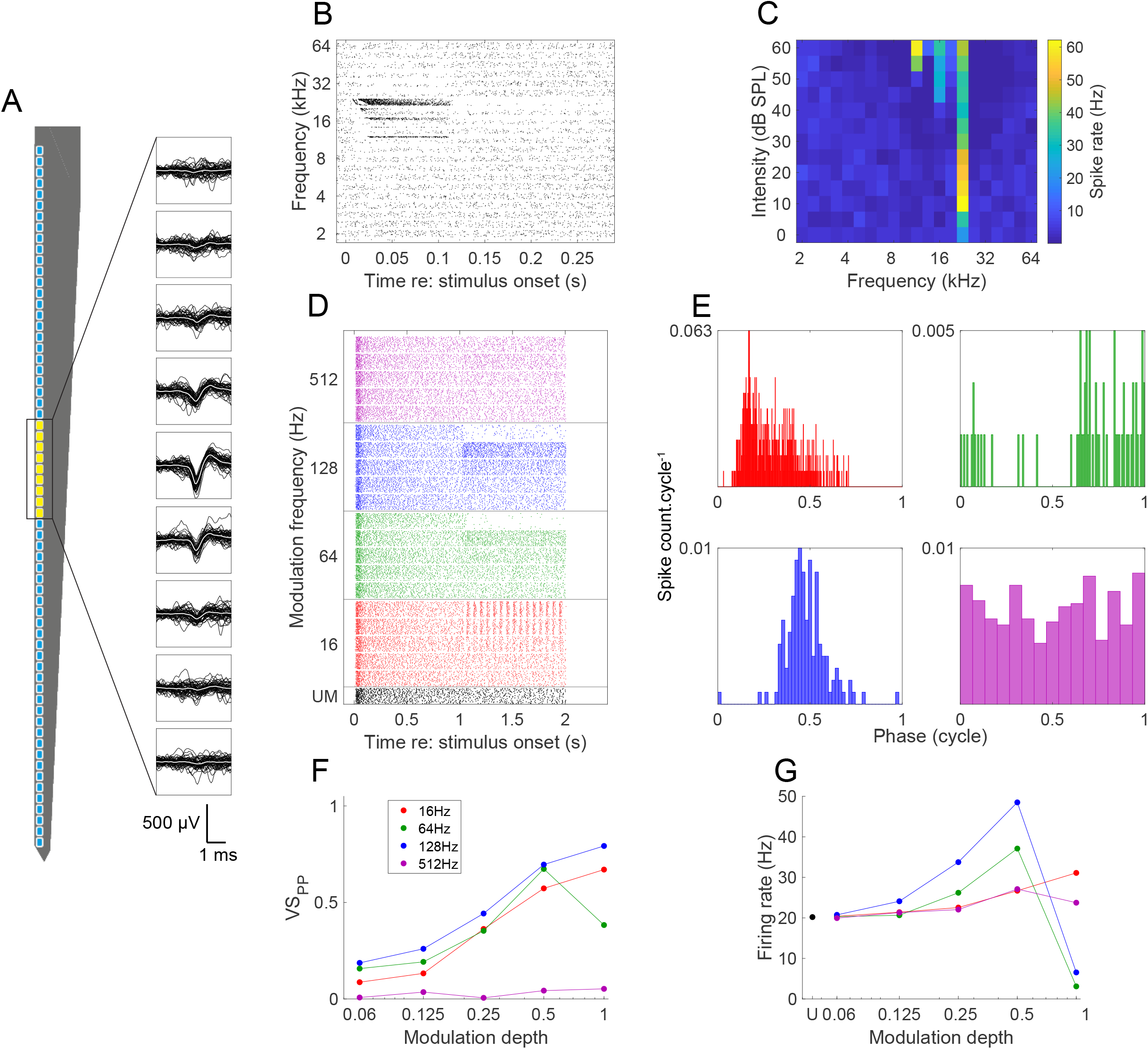
Example unit from the ICc with sustained responses. (A) Spike waveforms detected on adjacent channels. (B) Dot raster plot of the pure tone responses with the ordinate sorted by frequency followed by intensity. (C) FRA of pure tone responses showing sharp tuning with a CF of 22.6 kHz and a minimum threshold of 0 dB. (D) Dot raster plot of responses to AM stimuli. Trials organized by decreasing modulation depth for the different modulation frequencies. Transition from UM to AM occurred at 1 s. (E) Cycle histograms show phase-locking ability for full modulation stimuli. Color scheme of modulation frequency matches that of D. Bin size is 0.13 ms. (F) Temporal coding of AM was quantified by calculating VS_PP_ for the AM period. (G) The average firing rate during the AM period served as a measure for rate coding to AM, and shows a non-monotonic relationship with modulation depth for 64 and 128 Hz modulation frequencies. All AM data shown are for a carrier noise intensity of 60 dB SPL.

When presented with broadband noise, the same unit responded to the sound onset with a well-timed single spike response with strong inhibition for the remainder of the UM noise stimulus (Figure 4D). Upon transition to the AM stimulus (at t = 1 s), a pronounced increase in firing was observed, but mainly at the higher modulation depths. Sharp peaks in the cycle histograms exemplify strong phase-locking at 16 Hz (red), 64 Hz (green) and 128 Hz (blue; Figure 4E). Even at 512 Hz, the firing rate co-modulated with the stimulus period at large modulation depths. VS_PP_ showed a progressive increase for modulation frequencies 16, 64 and 128 Hz, up to a remarkable precision of >0.9 at full modulation depth (*m* = 1; Figure 4F). The VS_PP_ at 512 Hz also increased to about 0.4 at a modulation depth of 0.5, but decreased at full modulation depth. In this unit the firing rate also depended strongly on the modulation depth of the AM stimulus (Figure 4G).

Figure 5 illustrates an ICc unit with a less well-timed response to AM stimuli and a non-monotonic relationship between firing rate and modulation depth. The unit was sharply tuned, and highly sensitive at its CF of 22.6 kHz, with an estimated minimum threshold of 0 dB, the lowest intensity that was presented (Figure 5C). At higher intensities the spectral sensitivity broadened. At CF, tone responses showed the shortest onset delay, and firing was sustained during the tone presentation. Following sustained firing during the tone, offset inhibition was observed. Inhibitory sidebands were also present, especially at frequencies above CF (Figure 5B,C).

The unit showed a sustained response to broadband noise with a well-timed onset (Figure 5D). Firing rate generally increased with modulation depth at low- to mid-range modulation frequencies. However, the relationship between firing rate and modulation depth was non-monotonic at modulation frequencies of 64 and 128 Hz, where firing rate initially increased, but then decreased abruptly with increasing modulation depth (Figure 5D,G). A non-monotonic relationship between firing rate and modulation depth was quite common among the recorded units. The decrease in firing rate at 64 Hz was so large that the VS_PP_ became very low as well owing to the many trials without a single spike (Figure 5D, E). This artificial decrease in VS_PP_ was not observed at 128 Hz, where the decrease in firing rate was less than at 64 Hz.

The units illustrated in Figure 4 and 5 thus illustrate the well-known, large functional heterogeneity of IC neurons (Palmer et al., 2013). The large diversity of FRAs is in agreement with previous results (Ehret et al., 2005). We did not attempt to classify the different FRAs, as previous clustering attempts were unsuccessful (Hernández et al., 2005; Palmer et al., 2013).

Figure 6 illustrates the temporal (Figure 6A-C) and rate response (Figure 6D-F) of the population of IC neurons to AM stimuli. IC neurons generally showed good phase-locking ability (as measured by VS_PP_) at 16, 64 and 128 Hz, but performed poorly at 512 Hz (Figure 6A). Since each stimulus repetition yields a VS_PP_ value, we could apply ROC analysis to convert both VS_PP_ and firing rate to d’. AM detection thresholds were calculated analogously to the behavioral detection thresholds, using a d’ of 1 as the cutoff. The population mean curves of d’ calculated from the VS_PP_ (d’_temporal_; Figure 6B) closely resembled the population mean curves of VS_PP_ themselves (Figure 6A). VS_PP_ on average increased with increasing modulation depths, except in the case of 64 and 128 Hz where VS_PP_ decreased at the highest modulation depths (i.e. non-monotonic pattern). This pattern was also reflected in the detection thresholds. A detection threshold to a modulation frequency of 16 Hz could be established in 79% of the units (Figure 6C). This proportion decreased with increasing modulation frequency, with 63% and 51% of units showing detectable thresholds for 64 and 128 Hz, respectively. This threshold was on average at a lower modulation depth than for 16 Hz, reflecting the overall better phase-locking at these frequencies. In contrast, only 6% of the population showed a threshold for AM at 512 Hz, illustrating the generally poor phase-locking at this modulation frequency. This suggested that temporal coding could not explain the good behavioral detection threshold at 512 Hz (Square in Figure 6C).

**Figure 6.**
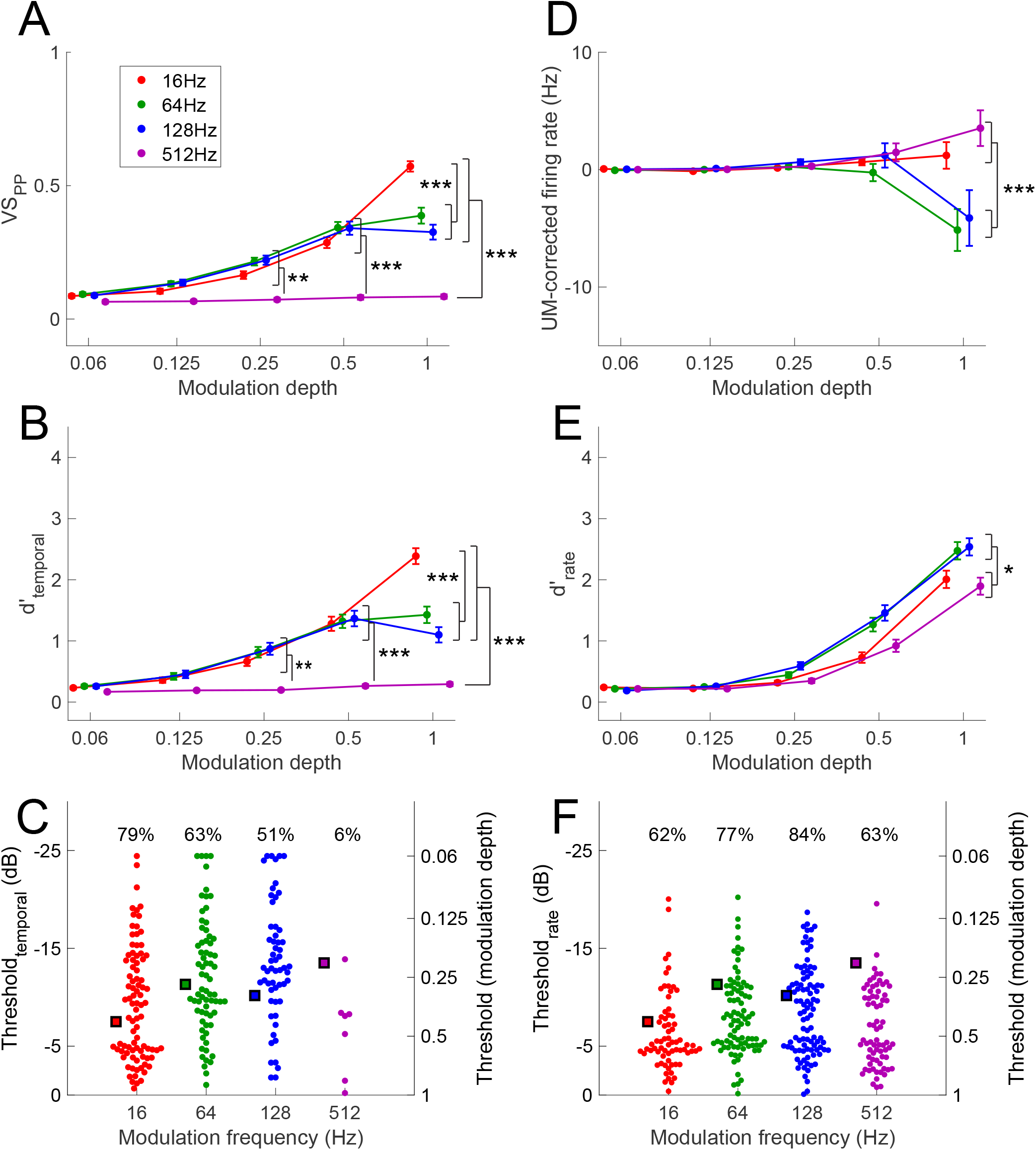
Population AM encoding. (A) Average relation between VS_PP_ and modulation depth. (B) Conversion of VS_PP_ to d’ using ROC analysis. (C) AM detection thresholds. Percentages denote the percentage of cells for which a threshold could be found. Thresholds were defined as the modulation depth for which d’ = 1. Black-edged markers indicate the mean detection thresholds obtained from behavioral experiments. (D) Average difference between firing rate of AM stimuli and that of the intensity-matched UM stimuli, as a function of modulation depth. (E-F) As B-C, but for firing rate instead of vector strength. Error bars in the figure show s.e.m.. * p<0.05, ** p<0.01, *** p<0.005 – as determined from two-way ANOVA and Tukey-Kramer post hoc test. All AM data shown are for a carrier noise intensity of 60 dB SPL.

The large variability in sound-evoked firing rates across IC neurons obscures the AM-evoked changes in the average firing rate. To better illustrate the effect of AM in the population data, we subtracted the firing rates evoked by the UM noise carrier at the same intensity from the AM-evoked firing rates before averaging (UM-corrected firing rate; Figure 6D). A small increase in the average firing rate can be seen for both 16 and 512 Hz at high modulation depths, whereas for both 64 and 128 Hz the firing rate on average showed a clear decrease at high modulation depths. All modulation frequencies showed an increase in the standard deviation of the AM firing rate as modulation depth increased, illustrating that both increases and decreases in firing rate were observed among different units. Note that the strong inhibition of firing rate in some units resulted in a decrease in mean VS_PP_ at these frequencies (Figure 6A), since a trial with no spikes is defined to have a trial VS of 0, which is ultimately averaged across repeated trials to obtain VS_PP_. Generally, this decrease in VS_PP_ therefore reflects the difficulty of assessing phase-locking when firing rate becomes very low.

Conversion of firing rate to d’ by ROC analysis created a more straightforward measure of performance (Figure 6E). At all four tested modulation frequencies, a gradual increase in d’ with increasing modulation depth was observed (Figure 6E). A measurable detection threshold was observed in most cases for the mid frequencies 64 and 128 Hz (77% and 84% respectively), whereas for 16 and 512 Hz this percentage was somewhat lower (62% and 63%, respectively). In the cells with a measurable threshold, these thresholds assumed similar values across frequencies (Figure 6F). In general, we found that rate-derived thresholds agree better with behavior.

As a control for potential effects of anesthesia, we also recorded a total of 99 units in awake, passively-listening animals (n = 4 mice). In awake and anesthetized animals similar results were obtained both with regard to phase-locking and firing frequencies at different modulation frequencies (Figure 7). Conversion of the neurometric measures of temporal (VS_PP_) and rate (firing rate) coding to d’ made it possible to compare the neural AM detection performance in the anesthetized and awake animals with the psychometric performance. At a modulation frequency of 16 Hz, behavioral performance was similar to the neurometric performance, both for temporal and for rate coding (Figure 7A and E, respectively). Both neurometric measures of AM coding showed similar trends as the psychometric measures at the midrange frequencies 64 Hz (Figure 7B, F) and 128 Hz (Figure 7C, G), with the exception of the drop in temporal coding performance seen at full modulation, as previously described. At 512 Hz, the d’ based on VS_PP_ was much smaller than the behavioral measure (Figure 7D), whereas the d’ based on firing rate was more similar (Figure 7H).

**Figure 7.**
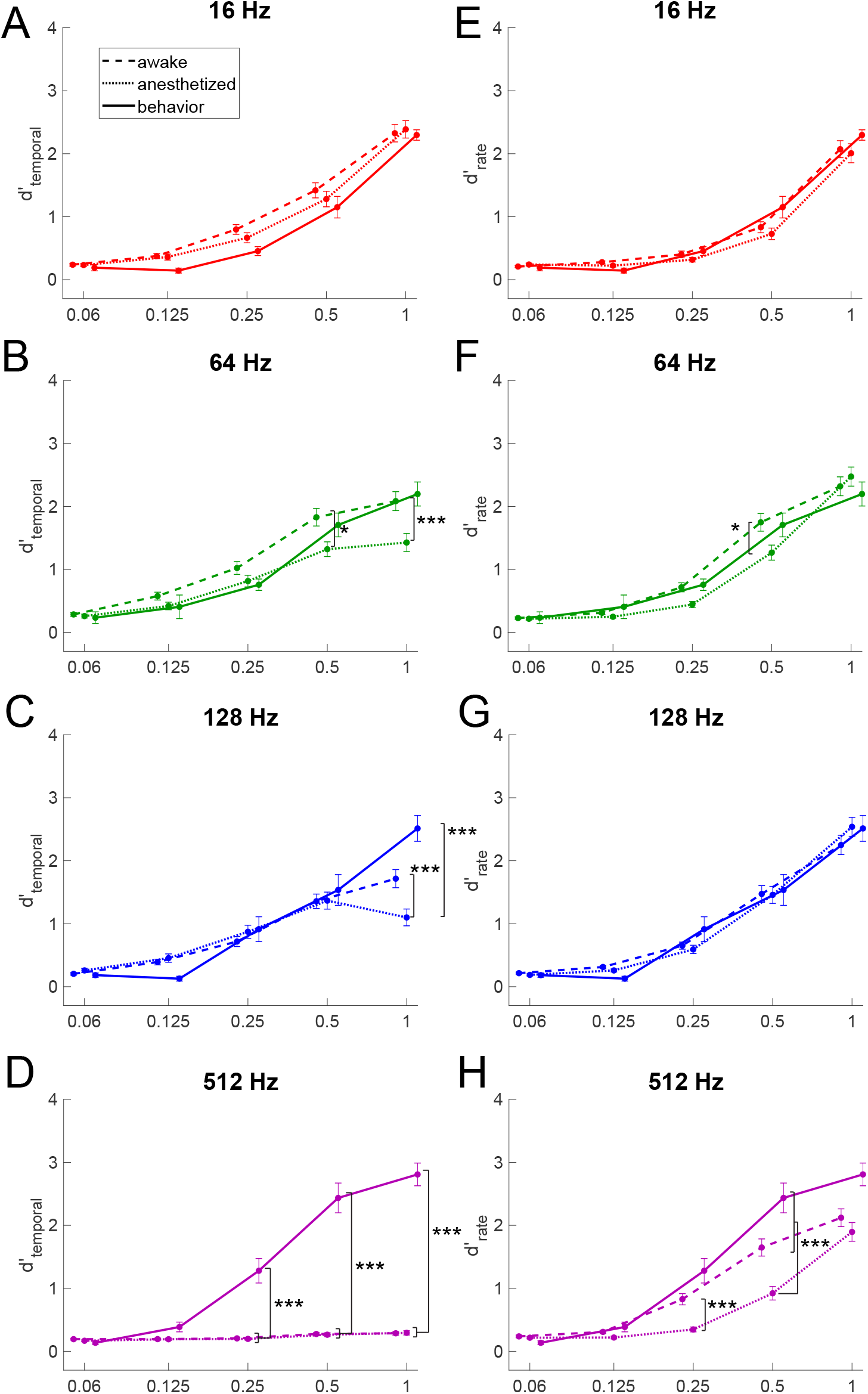
Comparing neurometric temporal and rate AM encoding of awake (dashed lines, n = 99 units) and anesthetized (dotted lines, n = 112 units) recordings to behavioral performance (solid lines, n = 8 animals). (A-D) VS_PP_-derived d’ plotted against modulation depth. (E-F) Firing rate-derived d’ plotted against modulation depth. mean ± s.e.m.. * p<0.05, *** p<0.005 – as determined from three-way ANOVA and Tukey-Kramer post hoc test. All AM data shown are for a carrier noise intensity of 60 dB SPL.

The overall performance of the awake units came closer to the behavioral performance. Especially at the mid frequencies, the relation between d’_temporal_ and modulation depth was less nonlinear (Figure 7B,C). At the highest modulation frequency, the d’_rate_ matched the behavioral d’ better than its counterpart in anesthetized animals (Figure 7H). However, d’_temporal_ at 512 Hz showed similarly low values as for the anesthetized animals (Figure 7D), indicating that phase-locking at the level of the IC unlikely plays a prominent role in encoding such high modulation frequencies. A comparison of thresholds between anaesthetized units, awake units and behavior is provided in Figure S3.

In order to investigate potential differences between the subdivisions of the IC in the encoding of AM stimuli, the recording electrodes were coated with the fluorescent dye DiO or DiI, and the probe tracks were reconstructed following postmortem histology. The location of individual units could be determined by aligning the histological sections to the Allen Brain Atlas CCFv3 (Wang et al., 2020) (Figure 8A). All single units obtained from anesthetized recordings (n = 112) and 82% (n = 81/99) of the units from awake recordings could thus be aligned to the reference atlas. The units originated from a large part of the IC except for its most lateral and caudal parts. The alignment also allowed us to assign each unit to a subnucleus within the IC based on the borders defined by the Allen Brain Atlas CCF. The proportion of units located within the central (ICc), dorsal (ICd) and external (ICe) subnuclei of the IC were 57%, 27% and 15%, respectively.

**Figure 8.**
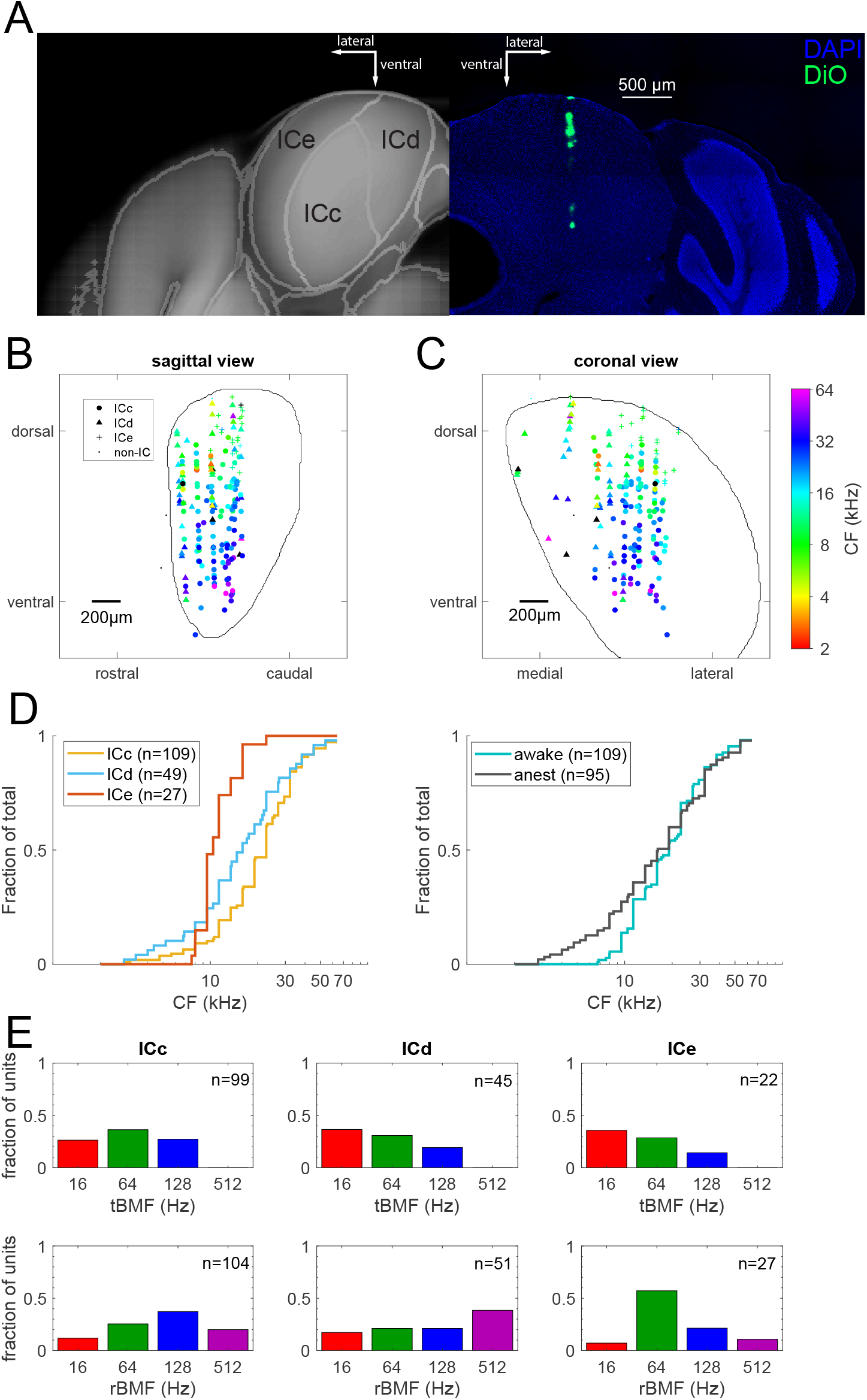
Spatial organization of pure tone and AM encoding of recorded units. Location of the units was obtained by registering probe tracts to the mouse Allen Brain Atlas. (A) Right: Coronal section showing the DiO-labelled probe tract. Left: mirror-image of corresponding virtual slice from the Allen Brain Atlas CCFv3 to illustrate the parcellation of the IC at the level of the probe tract. (B) Sagittal view of units in the left IC. Color of symbols represents CF and shape represents location of units within subnuclei of the IC. (C) as (B), but in coronal view. (D) Left: Cumulative distribution of CFs in the different subnuclei of the IC. Right: Cumulative distribution of CFs in recordings from anesthetized and awake animals. (E) Top: normalized distribution of temporal best modulation frequencies (tBMF) in the different subnuclei of the IC. Bottom: normalized distribution of rate best modulation frequencies (rBMF) within the different subnuclei of the IC. Chi-Squared tests showed significant differences among all the subnuclei in their rBMF distributions, but not for tBMFs.

**Figure 9.**
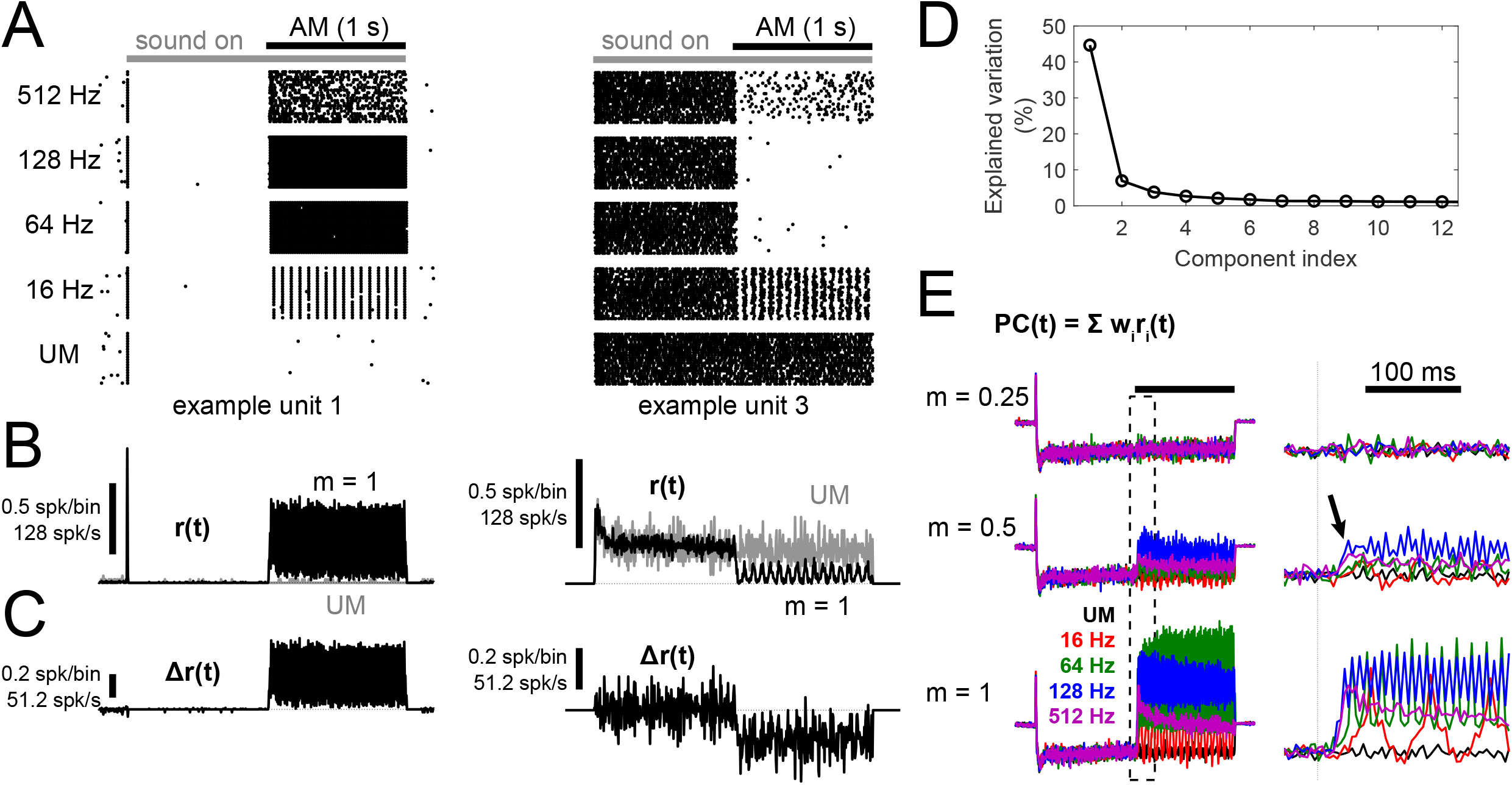
Principal component analysis of the difference in responses to AM or UM stimuli. (A) Dot raster plot of two example units in response to fully modulated stimuli at various frequencies and the unmodulated control stimulus (UM). (B) Average peristimulus time histograms (PSTHs) of the same units for all fully modulated stimuli (black; m = 1) and for the unmodulated control stimulus (gray). Bin size: 1/256 Hz ≈ 3.9 ms. (C) Difference between both PSTHs in B (PSTH_diff_), which was used as the input to the principal component analysis (PCA). (D) Scree plot showing the variation explained by each of the principal components in descending order. Only the first 12 components are shown. (E) Average amplitude of PC1 in response to stimuli of various modulation frequencies increased with modulation depth, showing the full stimulus (left) and zoomed in at the AM transition (right; dash box in left). Note the larger and earlier response to 512 and 128 Hz stimuli at m = 0.5 (arrow). Bin size: 1/256 Hz ≈ 3.9 ms.

Plotting the CF of the units versus their location revealed the dorsolateral to ventromedial tonotopic gradient of the IC (Portfors et al., 2011; Stiebler et al., 1985), in both a sagittal (Figure 8B) and a coronal view (Figure 8C). The distribution of CFs in ICc neurons spanned the entire frequency range of presented stimuli, but peaked between 16 and 32 kHz. Neurons sampled from the ICd had a similar distribution of CFs as in the ICc, while that of the ICe clustered between 8 and 16 kHz (Figure 8D left). The CF distribution of the neurons recorded under anesthesia and awake were generally similar (Figure 8D right).

We did not observe any obvious spatial clustering or gradient in AM encoding, as measured by the d’ thresholds calculated from VS_PP_ and firing rate (results not shown).

The preferred modulation frequencies were compared between the sampled neurons of the subregions of the IC. This was done for both their tBMF (Figure 8E top) and rBMF (Figure 8E bottom). Units in the ICc, ICd and ICe generally had tBMFs at one of the lower modulation frequencies. No significant difference was observed in the proportions of tBMFs among the three subregions when tested using a Chi-squared test (p=0.416). For the rBMF, the differences in proportion between all subregions were significant (p-value: ICd vs. ICe = 0.004, ICc vs. ICe = 0.018 & ICc vs. ICd = 0.038; Chi-squared test). In contrast to some previous studies (Baumann et al., 2011; Rodríguez et al., 2010; Schnupp et al., 2015; Schreiner et al., 1988), we did not find evidence for a gradient in tBMF or rBMF or for local clustering within the IC.

Our comparison of behavioral and neurometric thresholds showed better agreement for rate coding than for temporal coding. A limitation of the analysis above is that VS_pp_ and firing rates were calculated for up to the end of the AM period, while an animal engaged in the task often responded earlier with less available information. This may underly the difference between our behavioral thresholds and those reported by Cai et al. (2020) using a different task structure. To further explore how an animal would utilize spike information to perform our behavioral task, we performed a simulation combining dimensionality reduction with ideas from diffusion decision models (Ratcliff et al., 2016). Dimensionality reduction methods map high-dimensional data (e.g. activities of many neurons) into one or a few variables chosen to preserve relevant information from the data (Cunningham et al., 2014). This low-dimensional representation provides a link between neuronal activity and behavioral decisions, similar to mapping stimuli onto a single “decision axis” in Signal Detection Theory (Macmillan et al., 2004).

Since our mice were trained to distinguish AM sounds from unmodulated sounds regardless of their frequencies, we postulate that they must be relying on the difference between the neuronal activity evoked by the different modulation frequencies at the largest modulation depth (*m* = 1) and the neuronal activity evoked by the unmodulated sound (Figure 9A-C). We performed a principal component analysis (PCA) on this difference, and found that the first component (PC1) could already explain 44% of the variation with a sharp decrease to only 6.9% for the next component (Figure 9D). This suggested that this single component already captured most of the relevant information. The PCA results in a series of weights that, when multiplied with the firing rates of individual neurons, maps the population activities to a time-varying PC1 signal (see Methods). We further postulate that the size of PC1 is related to how well AM could be detected. Indeed, its amplitude showed a graded increase with increasing modulation depth across all frequencies (Figure 9E).

We were interested to learn more about the properties of neurons that best contributed to PC1. In Figure 10A we therefore sorted all neurons from anesthetized recordings according to their (absolute) weighting in PC1 and plotted their average response to noise onset and different AM frequencies as a heat map. The neuron illustrated in Figure 4 had the largest weight, whereas the neuron of Figure 5 came out halfway. Although the top 4 units showed an onset response to the broadband noise carrier (Figures 9A,B, 10A) and off-CF tones (Figure 4), onset response alone did not predict a good response to AM as demonstrated by the numerous onset units poorly represented in PC1. The next few units showed a mixture of sustained and build-up responses to broadband noise. Some lower ranking units were great phase-lockers, but showed little change in firing rate upon transition into AM. Figure 10B-C showed how the first 5 units encode AM in vector strength and firing rate, respectively.

**Figure 10.**
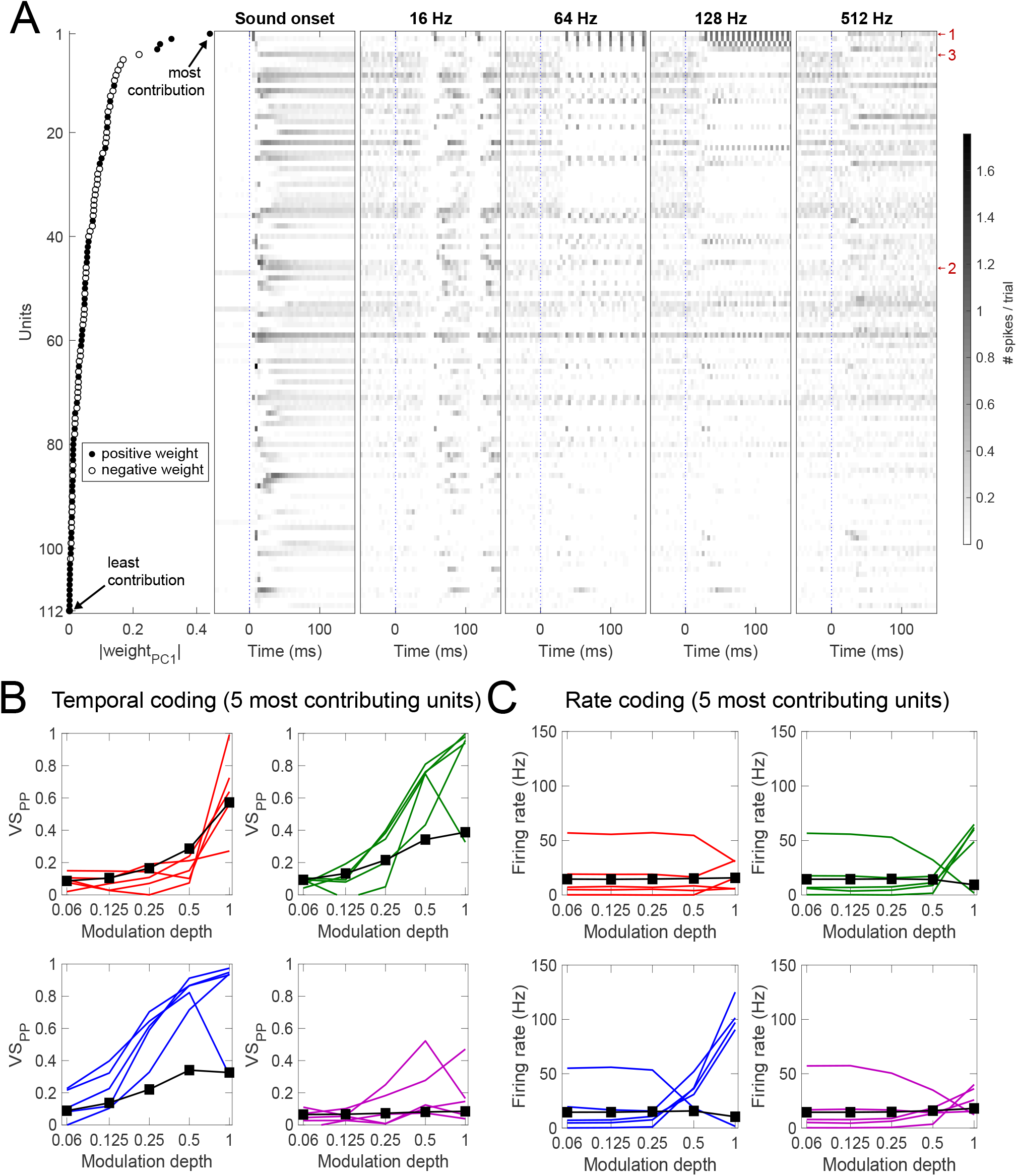
Overview of neurons from anaesthetized recordings ranked according to their contribution to PC1. (A) All neurons from anaesthetized recordings sorted by their contribution to PC1 (left panel, |weight_PC1_|), their average PSTH in response to sound onset (second from left) or to 16, 64, 128 or 512 Hz AM at full modulation. Closed and open symbols indicate positive and negative weights, respectively. Time is relative to either sound onset or transition to AM noise (blue, vertical dotted lines). Bin size: 1/256 Hz ≈ 3.9 ms. Small red arrows with numbers indicate example units. (1: Figure 4, 9A-C; 2: Figure 5; 3: Figure 9A-C) (B) VS_PP_ of top 5 units as a function of modulation frequency and depth. (C) Firing rate of top 5 units as a function of modulation frequency and depth. Modulation frequency color scheme in (B) and (C) follows that of previous figures: red = 16 Hz, green = 64 Hz, blue = 128 Hz and purple = 512 Hz.

In Figure 11 we did the same analysis for the units recorded during awake recordings. Top awake units showed a mixture of onset, sustained and build-up responses to broadband noise. Combining the awake and anesthetized recordings, the distinguishing feature of top neurons appears to be a strong change in firing rate upon AM (Figures 10B, 11B), although many of them are also good phase-lockers (Figures 10C, 11C). On the other hand, the vast majority of the neurons did show modulation in their firing rate to 16 Hz AM. In conclusion, the top contributors of PC1 generally had low spontaneous firing rates, strong and generally well-timed responses to AM, but showed a diverse (onset, adapted or sustained) response to the noise carrier.

**Figure 11.**
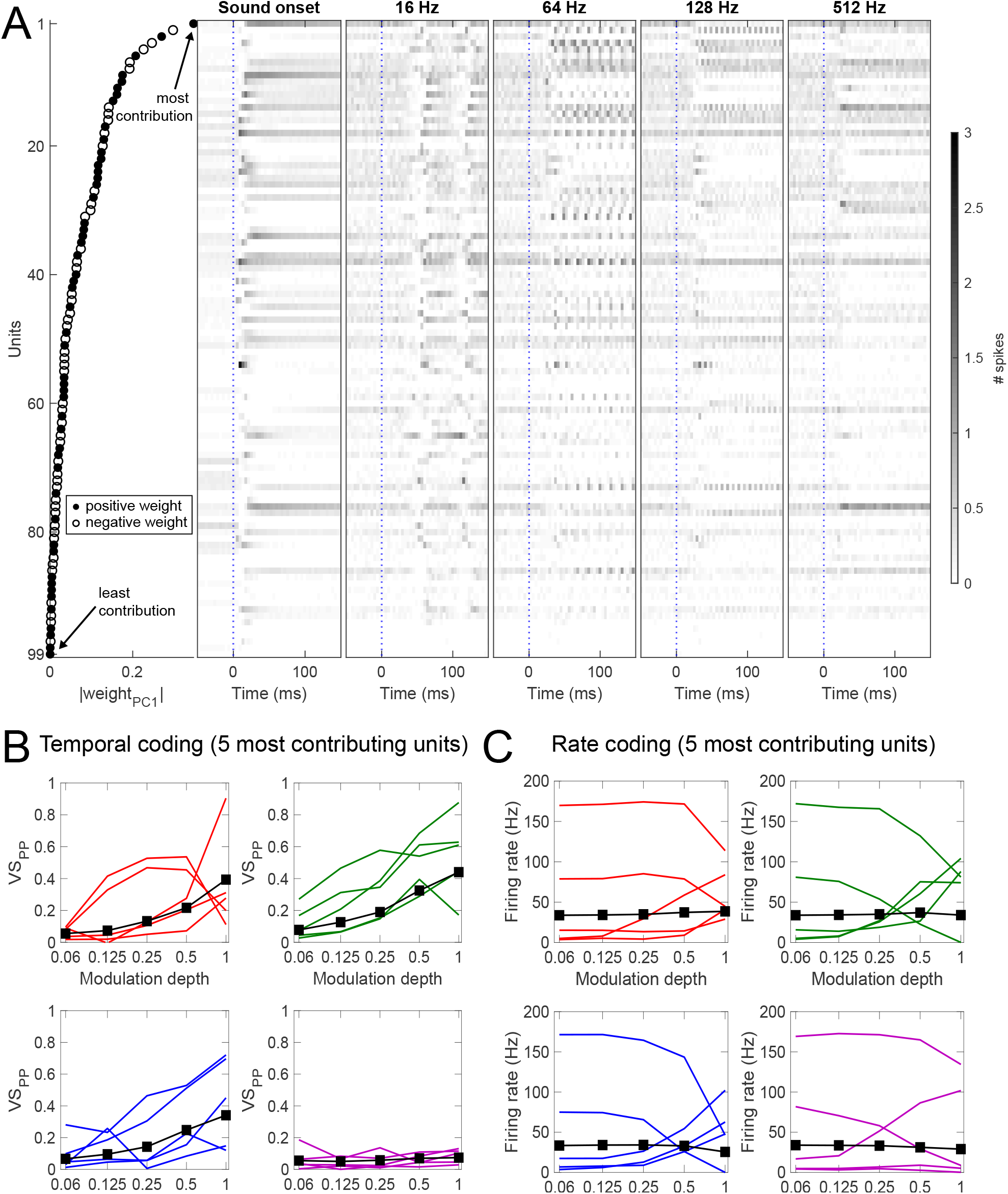
Overview of neurons from awake recordings ranked according to their contribution to PC1. (A-C) As Figure 10A-C, respectively, except recordings were from awake animals.

**Figure 12.**
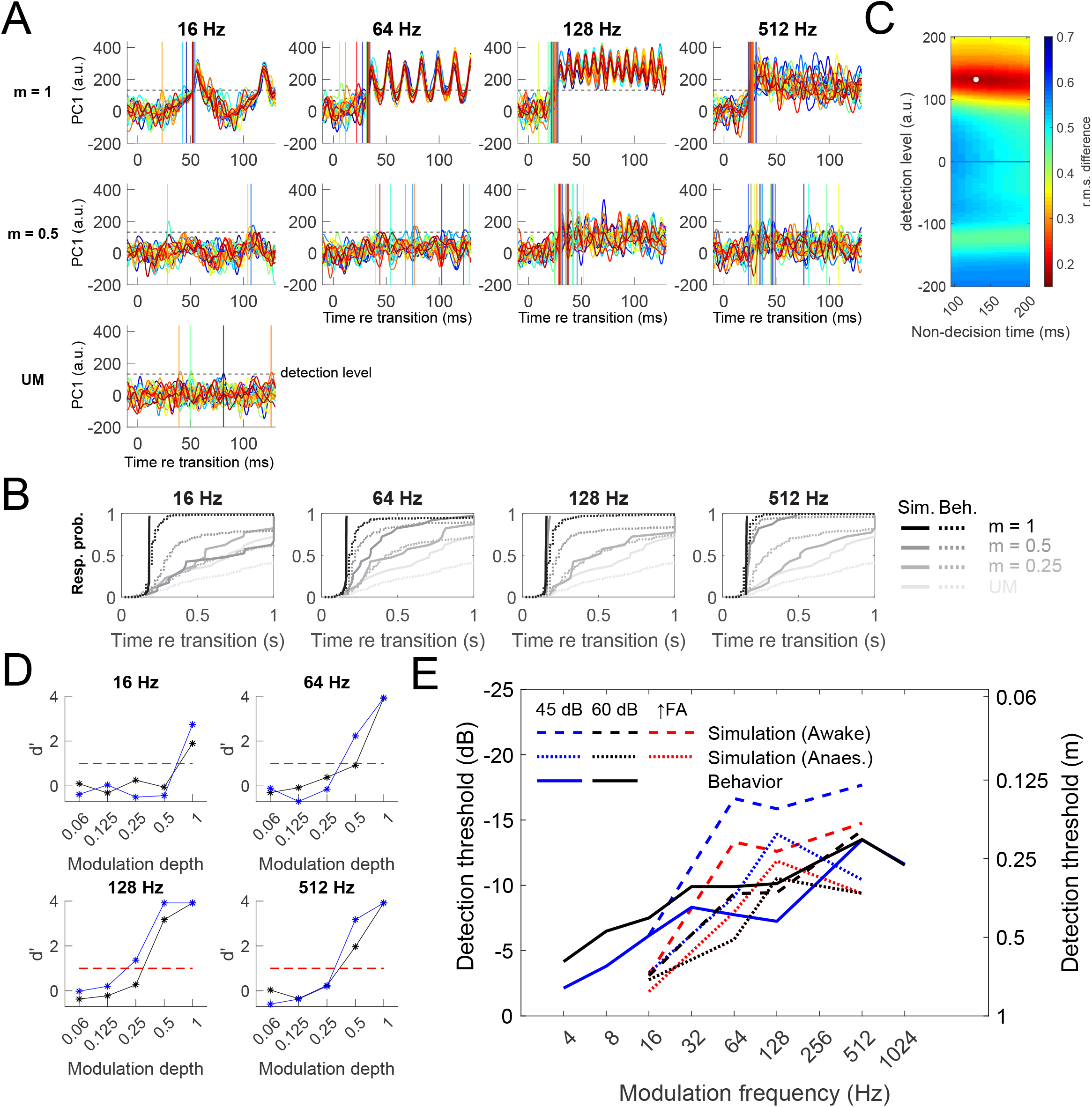
Using PCA of neuronal responses to AM stimuli to infer behavioral thresholds. (A) Single trial evolution of PC1 over time for selected modulation frequencies and depths at 60 dB for the data from the anaesthetized mice. Traces were created from kernel spike density functions with a Gaussian kernel (sigma: 2 ms). Horizontal lines represent the optimal “detection level” of the model that best fitted behavioral response latencies combined across all mice, and vertical lines represent the point at which PC1 exceeded this level, with colors corresponding to the PC traces. The model adds an additional non-decision time of 130 ms. (B) Comparison between cumulative distributions of simulated and behavioral response latency at different modulation frequencies and depths. (C) Root-mean-square error to response latency cumulative distributions as a function of parameters. White dot indicates location of the minima (non-decision time: 130 ms; detection level: 132 a.u.) (D) d’ calculated from simulated response latencies for 45 dB (blue) and 60 dB (black). (E) AM detection thresholds derived from the model compared to those from behavior. Red broken lines represent simulation thresholds with increased false alarm rate for 45 dB.

Whereas we performed the PCA with the trial-averaged response, during the behavioral task the animals only received information from single trials for each response decision. We further simulated this by projecting the single trial responses onto PC1 (Figure 12A), followed by measuring how long it took to cross a decision level at each trial. The decision level was obtained from a fit to the data. This threshold crossing readily captured the modulation frequency dependence in response latencies (Figure 3D) at full modulation depth (*m* = 1; Figure 12B). Interestingly, at a modulation depth of 0.5, the phase-locked fluctuation of PC1 at 16 Hz quite often did not reach the threshold, despite the vector strength being comparable to the 64 and 128 Hz stimuli (Figure 6A). It is possible that the longer modulation period at 16 Hz led to greater variability in spike time among neurons, making synchronous firing less likely.

Using this threshold crossing method, we simulated response latencies from single trial neuronal activities (Figure 12B-D), and subsequently estimate d’ and AM detection thresholds in the same way as the behavioral test (Figure 12D-E). The thresholds estimated from simulated detection latencies for the 60 dB stimuli captured the high pass pattern in the behavioral response, with 16 Hz being poorly distinguished from unmodulated noise, but progressively better discrimination for increasing modulation frequencies (Figure 12E). The sensitivity at 512 Hz modulation, however, did decrease slightly for the anaesthetized data, as was also apparent in the population firing rate d’ (Figure 6E). A slightly lower simulated false alarm rate at 45 dB (especially with awake data) led to an overestimation of AM sensitivity, which was corrected when false alarm rate at 60 dB was used instead to calculate *d’* (red broken lines in Figure 12E; see Discussion). Overall, we conclude that our model captured the modulation frequency dependence of the behavioral AM detection thresholds.

## 4. Discussion

We compared psychometric and neurometric detection of AM in the mouse using a new stimulus, which precluded spectral broadening as a cue. Their AM detection thresholds improved with modulation frequencies up to around 512 Hz, with comparatively poor performance at the lowest tested frequency (4 Hz) and very good performance at a high frequency (512 Hz). We compared these behavioral thresholds with neurometric thresholds based on the AM responses of single units in the IC. A PCA-based threshold method that took latencies into account did provide an adequate fit, especially in awake animals.

### 4.1. Excluding spectral broadening as a cue for AM detection

It has been assumed in AM studies early on that the spectrum of wideband noise do not change upon modulation (Viemeister, 1979). However, as we illustrated in Figure S1, AM of a noise carrier results in spectral broadening, which becomes substantial at the high modulation frequencies at which the mouse excelled. To ascertain that the mouse did not use these spectral cues, we compensated for the addition of sidebands, thus excluding a contribution of spectral (or intensity) cues to the measured AM detection thresholds. This method might be useful to retest the measured high-modulation-frequency sensitivities for other species, including humans.

### 4.2. Comparison of behavioral AM detection with earlier study in the mouse and with other species

The mice were poor at detecting AM at low modulation frequencies, but remarkably good at detecting high modulation frequencies. At 512 Hz, a comparison with other species indicated that the detection threshold of -13.5 dB placed the mice at a level similar to the best performing species (Carney et al., 2013; Dent et al., 2002; Henderson et al., 1984; Henry et al., 2016; Joris et al., 2004; Kelly et al., 2006). The relatively low thresholds at high modulation frequencies may enable mice to detect for example the fast amplitude modulations in brief ultrasonic vocalizations (Cai et al., 2020). Our study is in general agreement with a recent behavioral study in mice, in which a band-pass characteristic was also found for the behavioral detection (Cai et al., 2020), but we found higher thresholds overall. One potential explanation for the poor performance of our mice at low modulation frequencies could be the structure of the behavioral task, in which we presented the different modulation frequencies within the same session. The mice may have opted to rely on changes in perceptual features that are more salient for higher modulation frequencies (e.g. residual pitch over fluctuation; Joris et al., 2004). However, a study in which thresholds in macaques were compared for a block vs. randomized design did not show differences (O’Connor et al., 2011). A more plausible cause is that our mice were allowed to respond before the sound had finished. Shorter evaluation times were shown to impede detection for lower modulation frequencies (O’Connor et al., 2011).

### 4.3. Neural responses to AM

The responses to SAM noise generally matched earlier reports (Joris et al., 2004; Rees et al., 2005). In mice, responses to AM are generally heterogeneous (Geis et al., 2009; Ono et al., 2017; Walton et al., 2002). The absence of phase-locking at 512 Hz in most cells, which was in agreement with previous results using whole-cell recordings (Geis et al., 2009), indicates that there already must be some degrading of temporal information at the level of the IC, since in mice, phase-locking to modulation frequencies >500 Hz is still prominent at the level of the lower brainstem (Müller et al., 2019; Pauli-Magnus et al., 2007). In other species, phase-locking to frequencies up to 1200 Hz has been observed in the IC (Rees et al., 2005).

Our use of phase-projected vector strength avoided the spurious high vector strength values at very low spike rates encountered with traditional calculations (Yin et al., 2011). It also provided a trial-by-trial phase-locking metric convertible to a d’ value through ROC analysis. This allowed us to better compare between temporal and rate coding of AM sound, and to compare these neurometric measures to the psychometric measurements.

### 4.4. Effect of sound intensity

We found a slight improvement of behavioral AM detection at 60 dB SPL compared to 45 dB SPL. In other species the effects have not been very pronounced either. There was no clear effect of sound intensity on behavioral AM detection in parakeets or chinchillas (Dooling et al., 1981; Salvi et al., 1982). Owls and starlings and humans detect AM slightly better at higher sound intensities (Dent et al., 2002; Klump et al., 1991; Kohlrausch et al., 2000), similar to what was observed here, whereas macaques have slightly worse performance at higher sound intensities (Moody, 1994). The mechanisms underlying intensity-dependent changes are complex, and will depend among others on the tuning properties of the units that are differentially recruited (Rees et al., 1987; Wang et al., 2021).

### 4.5. Effect of anesthesia

Interestingly, the neuronal data obtained in awake animals approached the behavioral high-pass characteristics (Figure 7) better than the data obtained under anesthesia. This was also the case for the d’ of the VS or of the firing rate, suggesting that anesthesia may have had a depressing effect on neuronal responses. In previous studies, anesthesia had variable impact on the auditory system. Ketamine-xylazine may increase minimum threshold and interpeak latencies of ABRs in mice (van Looij et al., 2004). Only a limited number of studies addressed the impact on responses to SAM stimuli. In the rat medial geniculate body, urethane anesthesia did not significantly affect firing rate or VS (Cai et al., 2016). Only mild effects of pentobarbital plus ketamine anesthesia were observed on responses to AM stimuli in the gerbil IC (Ter-Mikaelian et al., 2007).

Relatively often we observed a strong decrease in firing rate at full modulation depth at the intermediate frequencies (64 and 128 Hz) after an initial increase with increasing modulation depth, similar to recordings in other rodents (Krebs et al., 2008; Zhang et al., 2003). In the rMTF, which is typically based on stimuli at full modulation depths, these would be band-reject neurons, whereas at low modulation depth these neurons showed a more band-pass-like character. In recordings from awake animals the decrease in firing at large modulation depths that was observed in many units seemed less conspicuous than in anesthetized animals. As this decrease in firing rate also may have contributed to the decrease in phase-locking, this decrease in non-linearity is likely to have contributed to the better match of the unanesthetized recordings with the behavioral data. Since well-timed, onset inhibition is likely to contribute to this nonmonotonicity (Geis et al., 2009), the effect of anesthesia may be to enhance this inhibitory component, but intracellular recordings would be needed to test this.

### 4.6. Neurometric vs. psychometric detection of AM

A main goal of our study was to try to understand which aspects of the neuronal responses within the IC informed the behavioral detection of AM. A coding scheme based on phase-locking already failed with the low VS at 512 Hz, as this frequency was behaviorally well detected. PSTH-based classification and many PCA-based methods include information that is not available to the animal when it responds early (Dong et al., 2013; Foffani et al., 2004; Malone et al., 2010).

We therefore opted for a two-step process in which spike responses of the whole stimulus were available to “train” the model to detect AM (PCA-step), whereas a threshold crossing enabled the model to respond at individual trials as soon as enough information was available. Since the animals were trained in a detection task, not to discriminate between different modulation frequencies, we performed PCA on the difference between the responses to the averaged AM and unmodulated stimuli, analogous to demixed PCA (Kobak et al., 2016). The PCA was performed on the neuronal dimension with time points as different “samples”. The first principal component (PC1) already explained >40% of the variance, allowing us to use only this PC1 to predict the behavioral responses from the IC population data. We used a linear dimensionality reduction method instead of newer, non-linear methods (McInnes et al., 2018; Moon et al., 2019; van der Maaten et al., 2008) for better interpretability.

We obtained simulated response delays with a simple level-crossing model based on the size of the population response mapped to PC1. Even though only two parameters needed to be estimated, our model overall provided a decent fit of the behavioral data, capturing the high-pass aspect of the behavioral detection. At 512 Hz, the large changes in firing rates of the top units ensured good performance. At 16 Hz, the predicted performance was much lower, despite decent phase-locking at these low frequencies. At low modulation frequencies, the slow amplitude changes lead to more dispersed spikes that sum less effectively across units, providing a possible explanation for the relatively poor behavioral detection.

Some limitations of the model were the following. While an animal performing the task may try to guess when a transition could occur based on past experience in trial timing, our simulated false positive responses were driven entirely by random fluctuations of the principal component, i.e. synchronous activity of the component units. This can be triggered by the temporal fine structure in the noise carrier or by co-modulation by non-auditory signals in the awake animal. Since we did not use frozen noise across different recordings, the lower firing rates at 45 dB will decrease crossings during catch trials, resulting in a “better” detection threshold. The lack of noise correlations across awake recording sessions may further lower false alarm rate at 45 dB. Recalculating the simulated threshold at 45 dB using the catch trials at 60 dB resulted in a better fit. An explicit false alarm rate independent of neuronal activity may therefore improve the model and better explain the underlying decision process of the animal. A further limitation is that neurometric and psychometric data were not collected simultaneously. The neuronal AM discriminability may improve in a task-engaged animal compared to our passively listening animals (Dong et al., 2013; Niwa et al., 2012a; Niwa et al., 2015; von Trapp et al., 2016).

Our interpretation of the results follows the lower envelope principle, which states that decisions to lick were based on a small number of the most sensitive neurons, whereas the many other, less sensitive neurons are largely disregarded (Barlow, 1972). It is also in line with the idea that the decision decoder that is responsible for the behavioral detection works optimally, since a characteristic of an optimal decoder is that it weighs sensory neurons by their sensitivity (Haefner et al., 2013; Jazayeri et al., 2006). A study in gerbils suggested that indiscriminate pooling, rather than a most-sensitive-neuron approach, best explains improved AM tone detection in adults relative to juveniles (Rosen et al., 2010). However, to more firmly establish causal relations and to discriminate between different decoder models, it would be necessary to assess the impact of manipulating the firing of IC neurons, for example using optogenetics (Marshel et al., 2019) or juxtacellular stimulation (Tanke et al., 2018).

## Abbreviations^1^

^1^ABR: auditory brainstem response
AM: amplitude modulation / amplitude-modulated
AUC: area under curve
d’: sensitivity index
IC: inferior colliculus
PSTH: peristimulus time histogram
PCA: principal component analysis
rBMF: rate best modulation frequency
rMTF: rate modulation transfer function
ROC: receiver operator characteristics
SAM: sinusoidal amplitude modulation / sinusoidally amplitude-modulated
tBMF: temporal best modulation frequency
tMTF: temporal modulation transfer function
UM: unmodulated
VS_PP_: phase-projected vector strength

## Acknowledgements

We thank Marcel van der Heijden for the concepts of the new AM stimulus, Peter Bremen for help with pilot multielectrode recording experiments, Jean Slenter for performing part of the mouse training, Jean Slenter, Sander Kruithof and Erica Sabel-Goedknegt for support with histology.

## References

Azeredo da Silveira R., Rieke, F. 2021. The geometry of information coding in correlated neural populations. Annu Rev Neurosci 44, 403–424.

Barlow, H.B. 1972. Single units and sensation: a neuron doctrine for perceptual psychology? Perception 1, 371–94.

Bartolo, R., Saunders, R.C., Mitz, A.R., Averbeck, B.B. 2020. Information-limiting correlations in large neural populations. J Neurosci 40, 1668–1678.

Baumann, S., Griffiths, T.D., Sun, L., Petkov, C.I., Thiele, A., Rees, A. 2011. Orthogonal representation of sound dimensions in the primate midbrain. Nat Neurosci 14, 423–5.

Bendor, D., Wang, X. 2007. Differential neural coding of acoustic flutter within primate auditory cortex. Nat Neurosci 10, 763–71.

Bremen, P., Massoudi, R., Van Wanrooij, M.M., Van Opstal, A.J. 2017. Audio-visual integration in a redundant target paradigm: a comparison between rhesus macaque and man. Front Syst Neurosci 11, 89.

Cai, H., Dent, M.L. 2020. Best sensitivity of temporal modulation transfer functions in laboratory mice matches the amplitude modulation embedded in vocalizations. J Acoust Soc Am 147, 337.

Cai, R., Richardson, B.D., Caspary, D.M. 2016. Responses to predictable versus random temporally complex stimuli from single units in auditory thalamus: impact of aging and anesthesia. J Neurosci 36, 10696–10706.

Carney, L.H., Ketterer, A.D., Abrams, K.S., Schwarz, D.M., Idrobo, F. 2013. Detection thresholds for amplitude modulations of tones in budgerigar, rabbit, and human. Adv Exp Med Biol 787, 391–8.

Carney, L.H., Zilany, M.S.A., Huang, N.J., Abrams, K.S., Idrobo, F. 2014. Suboptimal use of neural information in a mammalian auditory system. J Neurosci 34, 1306–13.

Cunningham, J.P., Yu, B.M. 2014. Dimensionality reduction for large-scale neural recordings. Nat Neurosci 17, 1500–9.

de Lafuente, V., Romo, R. 2006. Neural correlate of subjective sensory experience gradually builds up across cortical areas. Proc Natl Acad Sci U S A 103, 14266–71.

Dent, M.L., Klump, G.M., Schwenzfeier, C. 2002. Temporal modulation transfer functions in the barn owl (Tyto alba). J Comp Physiol A Neuroethol Sens Neural Behav Physiol 187, 937–43.

Dong, C., Qin, L., Liu, Y., Zhang, X., Sato, Y. 2011. Neural responses in the primary auditory cortex of freely behaving cats while discriminating fast and slow click-trains. PLoS One 6, e25895.

Dong, C., Qin, L., Zhao, Z., Zhong, R., Sato, Y. 2013. Behavioral modulation of neural encoding of click-trains in the primary and nonprimary auditory cortex of cats. J Neurosci 33, 13126–37.

Dooling, R.J., Searcy, M.H. 1981. Amplitude-modulation thresholds for the parakeet (Melopsittacus undulatus). Journal of Comparative Physiology 143, 383–388.

Ehret, G., Schreiner, C. 2005. Spectral and intensity coding in the auditory midbrain. In: Winer, J., Schreiner, C., (Eds.), The inferior colliculus. Springer, New York. pp. 312–345.

Foffani, G., Moxon, K.A. 2004. PSTH-based classification of sensory stimuli using ensembles of single neurons. J Neurosci Methods 135, 107–20.

Geis, H.-R., Borst, J.G.G. 2009. Intracellular responses of neurons in the mouse inferior colliculus to sinusoidal amplitude-modulated tones. J Neurophysiol 101, 2002–16.

Geis, H.R.A.P., van der Heijden, M., Borst, J.G.G. 2011. Subcortical input heterogeneity in the mouse inferior colliculus. J Physiol 589, 3955–67.

Goris, R.L.T., Ziemba, C.M., Stine, G.M., Simoncelli, E.P., Movshon, J.A. 2017. Dissociation of choice formation and choice-correlated activity in macaque visual cortex. J Neurosci 37, 5195–5203.

Guo, Z.V., Li, N., Huber, D., Ophir, E., Gutnisky, D., Ting, J.T., Feng, G., Svoboda, K. 2014. Flow of cortical activity underlying a tactile decision in mice. Neuron 81, 179–94

Haefner, R.M., Gerwinn, S., Macke, J.H., Bethge, M. 2013. Inferring decoding strategies from choice probabilities in the presence of correlated variability. Nat Neurosci 16, 235–42.

Henderson, D., Salvi, R., Pavek, G., Hamernik, R. 1984. Amplitude modulation thresholds in chinchillas with high-frequency hearing loss. J Acoust Soc Am 75, 1177–83.

Henry, K.S., Neilans, E.G., Abrams, K.S., Idrobo, F., Carney, L.H. 2016. Neural correlates of behavioral amplitude modulation sensitivity in the budgerigar midbrain. J Neurophysiol 115, 1905–16.

Hernández, O., Espinosa, N., Pérez-González, D., Malmierca, M.S. 2005. The inferior colliculus of the rat: a quantitative analysis of monaural frequency response areas. Neuroscience 132, 203–17.

Jazayeri, M., Movshon, J.A. 2006. Optimal representation of sensory information by neural populations. Nat Neurosci 9, 690–6.

Johnson, J.S., Yin, P., O’Connor, K.N., Sutter, M.L. 2012. Ability of primary auditory cortical neurons to detect amplitude modulation with rate and temporal codes: neurometric analysis. J Neurophysiol 107, 3325–41.

Joris, P.X., Schreiner, C.E., Rees, A. 2004. Neural processing of amplitude-modulated sounds. Physiol Rev 84, 541–77.

Kelly, J.B., Cooke, J.E., Gilbride, P.C., Mitchell, C., Zhang, H. 2006. Behavioral limits of auditory temporal resolution in the rat: amplitude modulation and duration discrimination. J Comp Psychol 120, 98–105.

Kettner, R.E., Thompson, R.F. 1985. Cochlear nucleus, inferior colliculus, and medial geniculate responses during the behavioral detection of threshold-level auditory stimuli in the rabbit. J Acoust Soc Am 77, 2111–27.

Klump, G.M., Okanoya, K. 1991. Temporal modulation transfer functions in the European starling (Sturnus vulgaris): I. Psychophysical modulation detection thresholds. Hear Res 52, 1–11.

Kobak, D., Brendel, W., Constantinidis, C., Feierstein, C.E., Kepecs, A., Mainen, Z.F., Qi, X.L., Romo, R., Uchida, N., Machens, C.K. 2016. Demixed principal component analysis of neural population data. Elife 5.

Kohlrausch, A., Fassel, R., Dau, T. 2000. The influence of carrier level and frequency on modulation and beat-detection thresholds for sinusoidal carriers. J Acoust Soc Am 108, 723–34.

Krebs, B., Lesica, N.A., Grothe, B. 2008. The representation of amplitude modulations in the mammalian auditory midbrain. J Neurophysiol 100, 1602–9.

Lu, T., Wang, X. 2004. Information content of auditory cortical responses to time-varying acoustic stimuli. J Neurophysiol 91, 301–13.

Macmillan, N.A., Creelman, C.D. 2004. Detection theory: A user’s guide Psychology press.

Malone, B.J., Scott, B.H., Semple, M.N. 2010. Temporal codes for amplitude contrast in auditory cortex. J Neurosci 30, 767–84.

Marshel, J.H., Kim, Y.S., Machado, T.A., Quirin, S., Benson, B., Kadmon, J., Raja, C., Chibukhchyan, A., Ramakrishnan, C., Inoue, M., Shane, J.C., McKnight, D.J., Yoshizawa, S., Kato, H.E., Ganguli, S., Deisseroth, K. 2019. Cortical layer-specific critical dynamics triggering perception. Science 365.

McInnes, L., Healy, J., Melville, J. 2018. UMAP: Uniform Manifold Approximation and Projection for Dimension Reduction. pp. arXiv:1802.03426.

Moody, D.B. 1994. Detection and discrimination of amplitude-modulated signals by macaque monkeys. J Acoust Soc Am 95, 3499–510.

Moon, K.R., van Dijk, D., Wang, Z., Gigante, S., Burkhardt, D.B., Chen, W.S., Yim, K., Elzen, A.V.D., Hirn, M.J., Coifman, R.R., Ivanova, N.B., Wolf, G., Krishnaswamy, S. 2019. Visualizing structure and transitions in high-dimensional biological data. Nat Biotechnol 37, 1482–1492.

Müller, M.K., Jovanovic, S., Keine, C., Radulovic, T., Rübsamen, R., Milenkovic, I. 2019. Functional development of principal neurons in the anteroventral cochlear nucleus extends beyond hearing onset. Front Cell Neurosci 13, 119.

Newsome, W.T., Britten, K.H., Movshon, J.A. 1989. Neuronal correlates of a perceptual decision. Nature 341, 52–4.

Niwa, M., Johnson, J.S., O’Connor, K.N., Sutter, M.L. 2012a. Active engagement improves primary auditory cortical neurons’ ability to discriminate temporal modulation. J Neurosci 32, 9323–34.

Niwa, M., Johnson, J.S., O’Connor, K.N., Sutter, M.L. 2012b. Activity related to perceptual judgment and action in primary auditory cortex. J Neurosci 32, 3193–210.

Niwa, M., O’Connor, K.N., Engall, E., Johnson, J.S., Sutter, M.L. 2015. Hierarchical effects of task engagement on amplitude modulation encoding in auditory cortex. J Neurophysiol 113, 307–27.

O’Connor, K.N., Johnson, J.S., Niwa, M., Noriega, N.C., Marshall, E.A., Sutter, M.L. 2011. Amplitude modulation detection as a function of modulation frequency and stimulus duration: comparisons between macaques and humans. Hear Res 277, 37–43.

Ono, M., Bishop, D.C., Oliver, D.L. 2017. Identified GABAergic and glutamatergic neurons in the mouse inferior colliculus share similar response properties. J Neurosci 37, 8952–8964.

Pachitariu, M., Steinmetz, N., Kadir, S., Carandini, M. 2016. Kilosort: realtime spike-sorting for extracellular electrophysiology with hundreds of channels. BioRxiv, 061481.

Palmer, A.R., Shackleton, T.M., Sumner, C.J., Zobay, O., Rees, A. 2013. Classification of frequency response areas in the inferior colliculus reveals continua not discrete classes. J Physiol 591, 4003–25.

Pauli-Magnus, D., Hoch, G., Strenzke, N., Anderson, S., Jentsch, T.J., Moser, T. 2007. Detection and differentiation of sensorineural hearing loss in mice using auditory steady-state responses and transient auditory brainstem responses. Neuroscience 149, 673–84.

Portfors, C.V., Mayko, Z.M., Jonson, K., Cha, G.F., Roberts, P.D. 2011. Spatial organization of receptive fields in the auditory midbrain of awake mouse. Neuroscience 193, 429–39.

Ratcliff, R., Smith, P.L., Brown, S.D., McKoon, G. 2016. Diffusion decision model: current issues and history. Trends Cogn Sci 20, 260–281.

Rees, A., Møller, A.R. 1987. Stimulus properties influencing the responses of inferior colliculus neurons to amplitude-modulated sounds. Hear Res 27, 129–43.

Rees, A., Langner, G. 2005. Temporal coding in the auditory midbrain. In: Winer, J., Schreiner, C., (Eds.), The inferior colliculus. Springer, New York. pp. 346–376.

Renart, A., de la Rocha, J., Bartho, P., Hollender, L., Parga, N., Reyes, A., Harris, K.D. 2010. The asynchronous state in cortical circuits. Science 327, 587–90.

Rodríguez, F.A., Read, H.L., Escabí, M.A. 2010. Spectral and temporal modulation tradeoff in the inferior colliculus. J Neurophysiol 103, 887–903.

Rosen, M.J., Semple, M.N., Sanes, D.H. 2010. Exploiting development to evaluate auditory encoding of amplitude modulation. J Neurosci 30, 15509–20.

Salvi, R.J., Giraudi, D.M., Henderson, D., Hamernik, R.P. 1982. Detection of sinusoidally amplitude modulated noise by the chinchilla. J Acoust Soc Am 71, 424–9.

Schnupp, J.W.H., Garcia-Lazaro, J.A., Lesica, N.A. 2015. Periodotopy in the gerbil inferior colliculus: local clustering rather than a gradient map. Front Neural Circuits 9, 37.

Schnupp, J.W.H., Hall, T.M., Kokelaar, R.F., Ahmed, B. 2006. Plasticity of temporal pattern codes for vocalization stimuli in primary auditory cortex. J Neurosci 26, 4785–95.

Schreiner, C.E., Langner, G. 1988. Periodicity coding in the inferior colliculus of the cat. II. Topographical organization. J Neurophysiol 60, 1823–40.

Shamash, P., Carandini, M., Harris, K., Steinmetz, N. 2018. A tool for analyzing electrode tracks from slice histology. bioRxiv, 447995.

Stiebler, I., Ehret, G. 1985. Inferior colliculus of the house mouse. I. A quantitative study of tonotopic organization, frequency representation, and tone-threshold distribution. J Comp Neurol 238, 65–76.

Stringer, C., Michaelos, M., Tsyboulski, D., Lindo, S.E., Pachitariu, M. 2021. High-precision coding in visual cortex. Cell 184, 2767–2778 e15.

Tanke, N., Borst, J.G.G., Houweling, A.R. 2018. Single-cell stimulation in barrel cortex influences psychophysical detection performance. J Neurosci 38, 2057–2068.

Ter-Mikaelian, M., Sanes, D.H., Semple, M.N. 2007. Transformation of temporal properties between auditory midbrain and cortex in the awake Mongolian gerbil. J Neurosci 27, 6091–102.

van der Maaten, L., Hinton, G. 2008. Visualizing Data using t-SNE. J Mach Learn Res 9, 2579–2605.

van Looij, M.A.J., Liem, S.S., van der Burg, H., van der Wees, J., De Zeeuw, C.I., van Zanten, B.G.A. 2004. Impact of conventional anesthesia on auditory brainstem responses in mice. Hear Res 193, 75–82.

Viemeister, N.F. 1979. Temporal modulation transfer functions based upon modulation thresholds. J Acoust Soc Am 66, 1364–80.

von Trapp, G., Buran, B.N., Sen, K., Semple, M.N., Sanes, D.H. 2016. A decline in response variability improves neural signal detection during auditory task performance. J Neurosci 36, 11097–11106.

Walker, K.M.M., Ahmed, B., Schnupp, J.W.H. 2008. Linking cortical spike pattern codes to auditory perception. J Cogn Neurosci 20, 135–52.

Walton, J.P., Simon, H., Frisina, R.D. 2002. Age-related alterations in the neural coding of envelope periodicities. J Neurophysiol 88, 565–78.

Wang, L., Narayan, R., Graña, G., Shamir, M., Sen, K. 2007. Cortical discrimination of complex natural stimuli: can single neurons match behavior? J Neurosci 27, 582–9.

Wang, Q., Ding, S.L., Li, Y., Royall, J., Feng, D., Lesnar, P., Graddis, N., Naeemi, M., Facer, B., Ho, A., Dolbeare, T., Blanchard, B., Dee, N., Wakeman, W., Hirokawa, K.E., Szafer, A., Sunkin, S.M., Oh, S.W., Bernard, A., Phillips, J.W., Hawrylycz, M., Koch, C., Zeng, H., Harris, J.A., Ng, L. 2020. The Allen Mouse Brain Common Coordinate Framework: A 3D reference atlas. Cell 181, 936–953 e20.

Wang, Y., Abrams, K.S., Carney, L.H., Henry, K.S. 2021. Midbrain-level neural correlates of behavioral tone-in-noise detection: dependence on energy and envelope cues. J Neurosci 41, 7206–7223.

Yin, P., Johnson, J.S., O’Connor, K.N., Sutter, M.L. 2011. Coding of amplitude modulation in primary auditory cortex. J Neurophysiol 105, 582–600.

Zhang, H., Kelly, J.B. 2003. Glutamatergic and GABAergic regulation of neural responses in inferior colliculus to amplitude-modulated sounds. J Neurophysiol 90, 477–90.

