## Supplemental Figures and Notes for "Neuronal responses in mouse inferior colliculus correlate with behavioral detection of amplitude modulated sound"

by van den Berg et al.

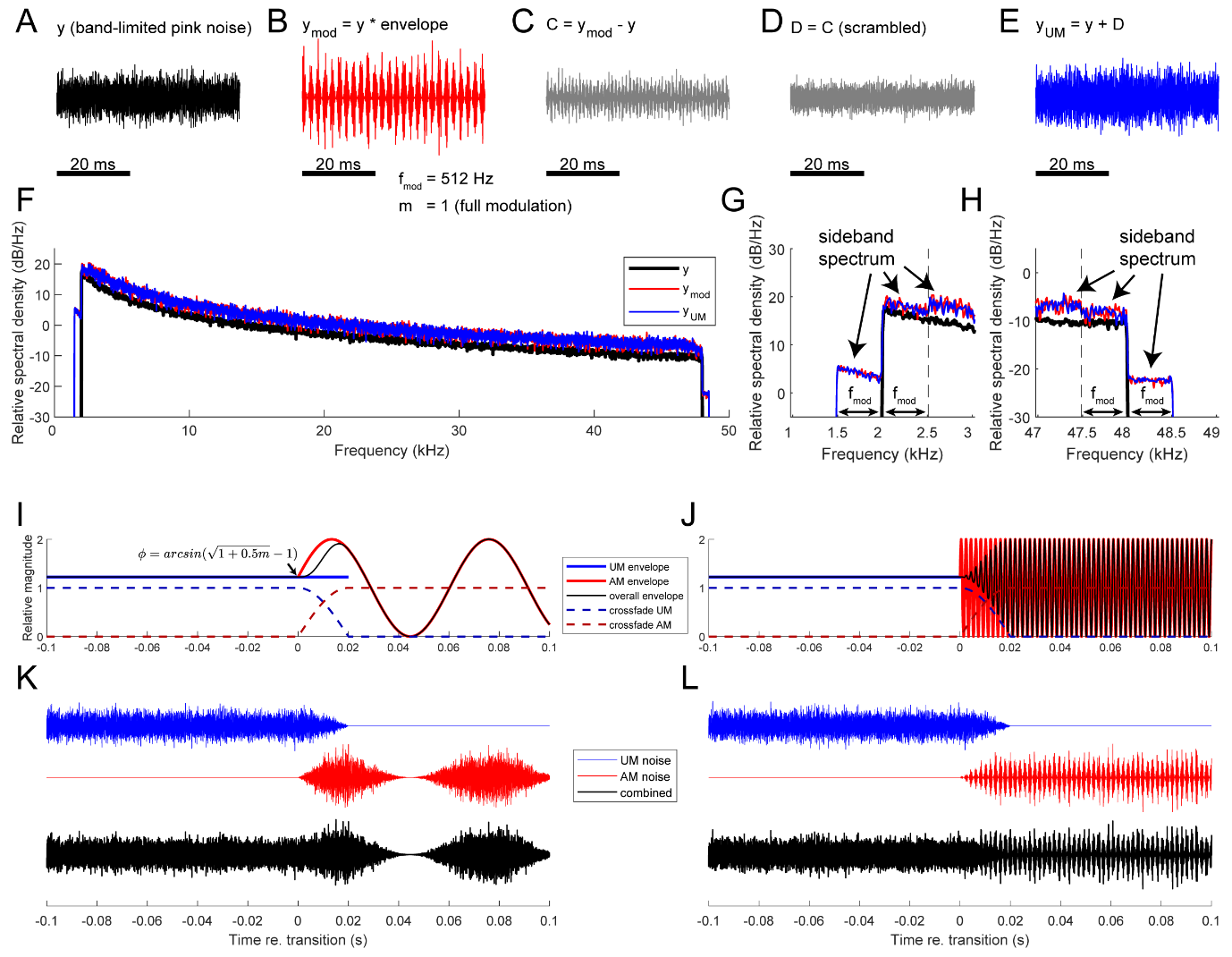

Figure S1 Generation of amplitude modulation stimulus. (A-E) Steps and respective waveforms of generating amplitude modulated and spectrally matched unmodulated noise. Band-limited pink noise (A) was modulated with an envelope $\left( 1+m\times sin \left( 2\pi f_{mod}t+\phi\right) \right)$ to generate a modulated noise, y_mod_ (B). In the example we used an $f_{mod}$ of 512 Hz and an $m$ of 1. The difference between the two are the modulation sidebands (C) of which we randomize the phase component to produce a “scrambled” version (D). These scrambled sidebands are added back to the original signal to produce a spectrally matched UM noise, y_UM_ (E). (F-H) Relative spectral density for the original pink noise from (A) (black; band-limited between 2-48 kHz), AM noise in (B) (red) and the UM noise in (E) (blue). Zooming into the lower- (G) and higher- (H) frequency edges reveal the spectral broadening and increase in overall intensity induced by the amplitude modulation (“sideband spectrum”, difference between red and black traces). The UM noise (blue traces) generated by our procedure is spectrally matched with the AM noise (red traces). (I-J) Intensity (solid lines) and cross-fading (broken lines) envelopes of the UM (blue) and AM (red) parts of a 16 Hz (I) and a 512 Hz (J) stimulus. Black lines indicate the overall intensity envelopes once the two parts are combined. (K-L) Example waveforms for crossfaded UM (blue), AM (red) and combined (black) noises for a 16 Hz (K) and 512 Hz (L) stimulus, respectively. The UM and AM parts are generated from independent instances of carrier noise.

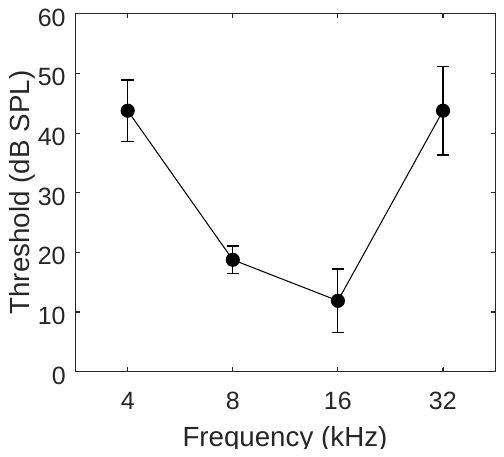

Figure S2. Hearing threshold of the mice. ABR thresholds (mean ± s.d.) of the B6CBAF1/JRj mice used in the behavioral experiments in this study (n = 8).

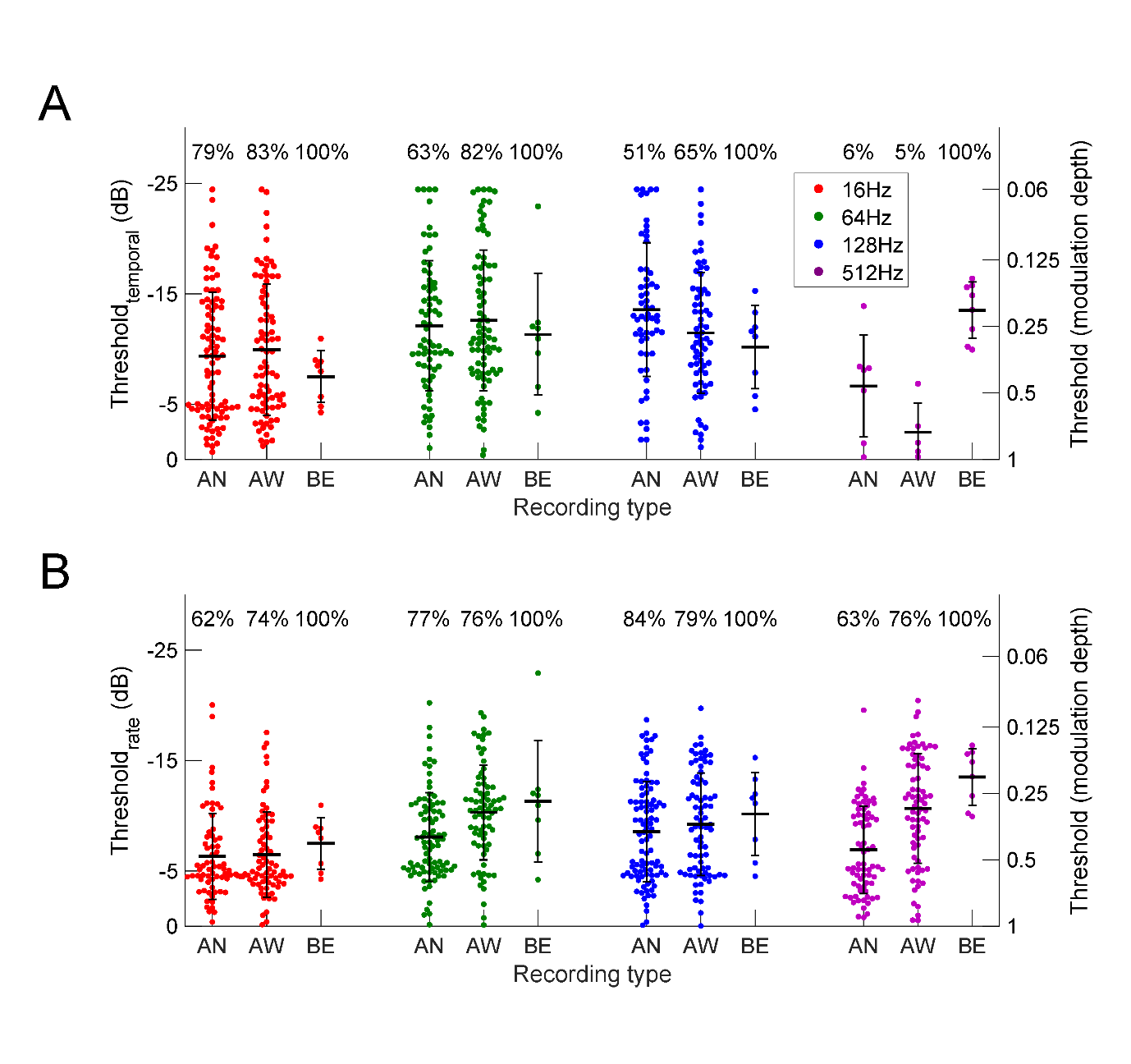

Figure S3 Comparison of AM detection thresholds from anesthetized, awake and behavioral recordings, as calculated from (A) VS-derived-d’ and (B) firing rate-derived-d’. Values above plotted thresholds show the percentage of units (or mice) that detected that modulation frequency. Horizontal black lines and error bars indicate the mean ± s.d. detection threshold for units (or mice) with detected thresholds. AN = anesthetized, AW = awake & BE = behavioral.

**Supplementary Note 1: Evaluation of effect of parameter choice on principal component analysis**

To test the robustness of the results of the principal component analysis (PCA) of neuronal responses, we systematically evaluated the effect of several parameter choices: temporal resolution (bin size), PSTH normalization, mean subtraction and inclusion of responses to different modulation depths.

Temporal resolution: bin sizes were varied between 0.98 ms (1/1024 Hz) to 500 ms (1/2 Hz).

PSTH normalization: SD of each unit’s PSTH was normalized to 1 or remained the same.

Inclusion of modulation depths: the difference in response is calculated either between the average response to all fully-modulated stimuli (m = 1) and the averaged response to unmodulated stimuli; or between the average response across all modulated stimuli (m = 0.06 to 1; “all mod”) and the averaged response to unmodulated stimuli.

Mean subtraction: the mean of the difference in response (*Δr_i_(t)*) for each unit was subtracted before input to PCA.

We evaluate these choices in two ways:

Firstly, how similar are the weights of the first principal component (PC1)? These weights indicate how much each neuron is contributing to PC1. The similarity of PC1s can be measured by the dot product between two PC1s; a dot product of 1 represents identical weights, and 0 represents highly dissimilar weights. Suppl. Figure S4A,C shows that the PCA result was very robust against choices of bin size and PSTH normalization. Normalization and mean-subtraction in general had little effect on the variation explained by PC1. The generally high value of the dot product indicates that the units were extracted largely in the same order to represent the PC1.

Secondly, how well can the principal component explain the difference in neuronal responses between UM and AM (i.e. *Δr_i_(t*); see section 2.9 ) As Figure S4B,D illustrates, including all modulation depths generally decreased explained variation (“m=1” traces vs “all m” traces). Most prominent, however, was the effect of temporal binning. This can be shown as a monotonic increase in variation explained by PC1 as bin size increases (upper left: Figure S4B and S4D). Larger bins, i.e. smoother data, allowed PC1 to explain more of the overall variability, presumably by averaging out temporal fluctuations in both noise and signal.

Despite differences in explained variation, the generally high dot product values indicated that all PC1s had largely similar weights for individual neurons. In the end, we chose not to perform normalization and mean-subtraction to avoid over-representation of low firing rate units, and to focus on the difference in firing rate. Un-normalized input also means that the resulting principal components remained in the firing rate scale.

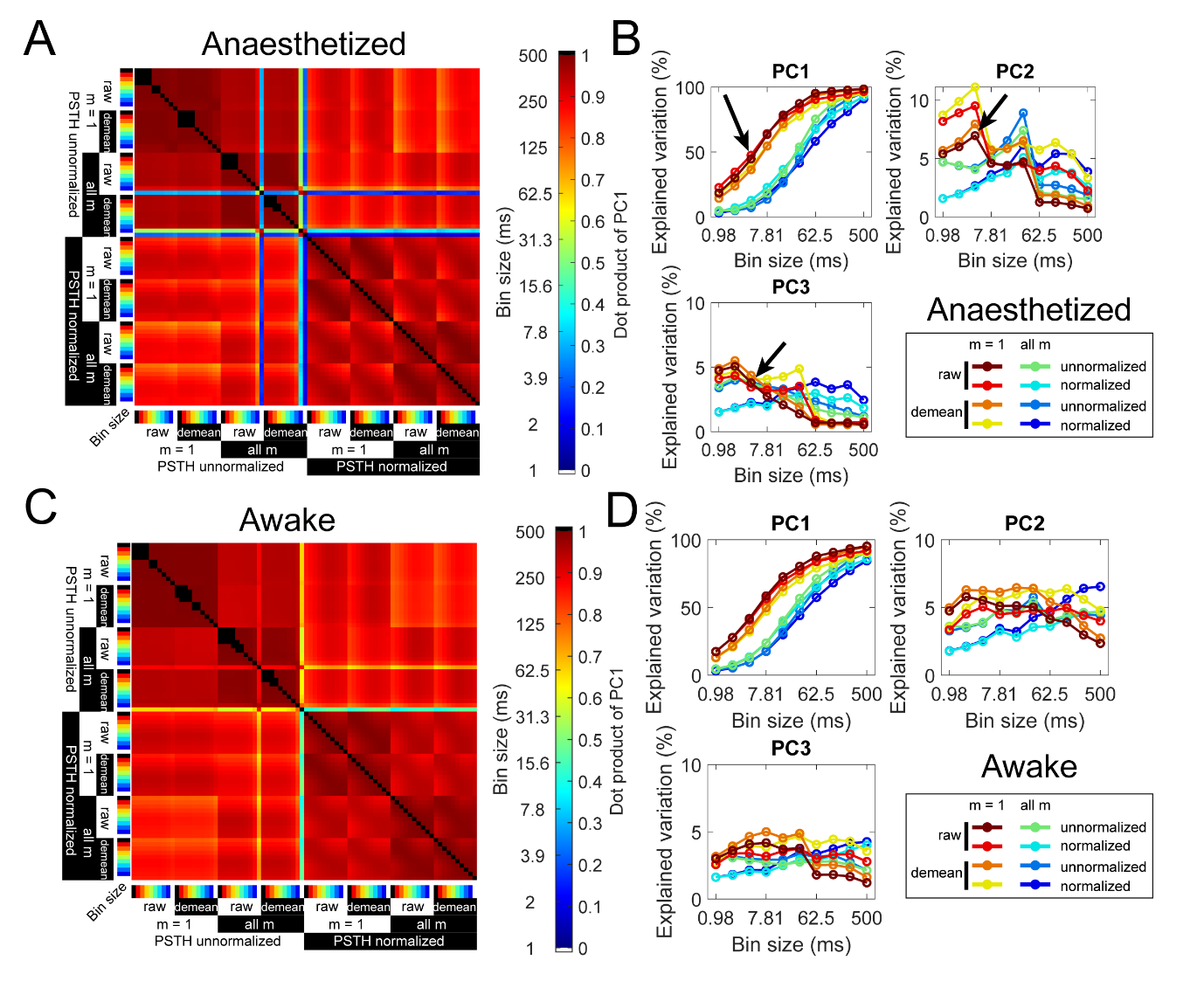

Figure S4: Robustness of PCA results against parameter choice in both anaesthetized and awake datasets. **(A)** Absolute values of dot products among PC1s from PCA using different PSTH bin sizes and parameters for anaesthetized dataset. The parameter choices were: whether the PSTH of each neuron was first normalized (‘normalized’ vs ‘unnormalized’), whether input was the difference between unmodulated response and the full modulation (‘m = 1’) or stimuli of all modulation depths (‘all m’), and whether the mean was subtracted from the PCA input (‘raw’ vs ‘demean’). **(B)** Explained variation plotted for the first three principal components (PC1-3) as a function of PSTH bin size and other parameter choices, as indicated in the boxes. Note the expanded y-scale for PC2 and PC3. Arrows indicate the parameter set shown in D-E and used in the main text. **(C)** Same as (A) but for the awake dataset. **(D)** Same as (B) but for the awake dataset.
